# Transcriptomic homogeneity and an age-dependent onset of hemoglobin expression characterize morphological PV types

**DOI:** 10.1101/2020.01.21.913103

**Authors:** Lin Que, David Lukacsovich, Csaba Földy

## Abstract

The diversity created by >100 different neural cell types fundamentally contributes to brain function and a central idea is that neuronal identity can be inferred from genetic information. Recent large-scale transcriptomic assays seem to confirm this hypothesis, but a lack of morphological information has limited the identification of several known cell types. For example, parvalbumin interneurons (PV-INs) comprise of a main transcriptomic cluster within all inhibitory cells. However, transcriptomics alone has not resolved the different morphological PV types that exist. To close this gap, we used single-cell RNA-seq in morphologically identified PV-INs, sampled from 10 days to 3 months-old mice and studied their transcriptomic states in the morphological, physiological, and developmental domains. Our findings reveal novel genes whose expression separately identify morphological types but corroborate an overall transcriptomic homogeneity among PV-INs. Surprisingly, morphological PV types display uniform cell adhesion molecule (CAM) profiles, suggesting that CAMs do not actively maintain their specificity of wiring after development. Finally, our results reveal a pronounced change of transcriptomic states between postnatal days 20 and 25, during which PV-INs display a rapid and unexpected onset of hemoglobin gene expression which remains stable in later development.

## Introduction

A central goal in brain research is the clear and comprehensive classification of cell types according to anatomical and physiological features, as well as by distinct molecular markers that unambiguously identify each type (Petilla Interneuron Nomenclature Group, 2008; Freund and Buzsáki, 1996; Klausberger and Somogyi, 2008; Zeng and Sanes, 2017; Booker and Vida, 2018). Recent single-cell RNA-seq assays have greatly facilitated classification efforts (e.g. Ziesel et al., 2015; Shekhar et al., 2016; Harris et al., 2018, Tasic et al., 2018) and increased expectations that transcriptomic information could explain the entirety of cell type-specific features, a crucial step towards understanding the multimodal identity of neural cell types. While several studies have employed single-cell RNA-seq to characterize physiological features (Cadwell et al., 2016; Földy et al., 2016; Fuzik et al., 2016; Muñoz-Manchado et al., 2018; Luo et al., 2019; Oláh et al., 2019; Winterer et al., 2019; Zheng et al., 2019), the relationship between transcriptomic information and morphology has not been investigated (Que et al., 2019).

PV-INs of the CA1 hippocampus are a particularly suitable model to study this problem, as they appear to be a physiologically homogenous population (Hu et al., 2014), yet have distinct morphological types. Morphologically, hippocampal PV-INs can be divided into 3 main cell types based on their axonal projections: axo-axonic cells (AAC) that specifically project to the axon initial segment of pyramidal cells; basket cells (BC), which establish synapses onto the perisomatic region of the postsynaptic neuron and thus restrict their axons to the pyramidal cell layer; and bistratified cells (BIC), whose axons target more distal dendrites and thereby extend their axons specifically in the oriens and radiatum (Booker and Vida, 2018; Maccaferri, 2005). Based on dendritic morphology, both basket and bistratified cells may each be further subdivided into horizontal (h) and vertical (v) subtypes (hBC, vBC; hBIC, vBIC, respectively; **Fig. 1**). Although these clear anatomical prototypes have been used to classify PV-INs, a certain continuity between morphological PV types is presumed to exist, where cells may display overlap of multiple morphological characteristics of the different types (Kohus et al., 2016; Booker and Vida, 2018).

**Fig. 1.**
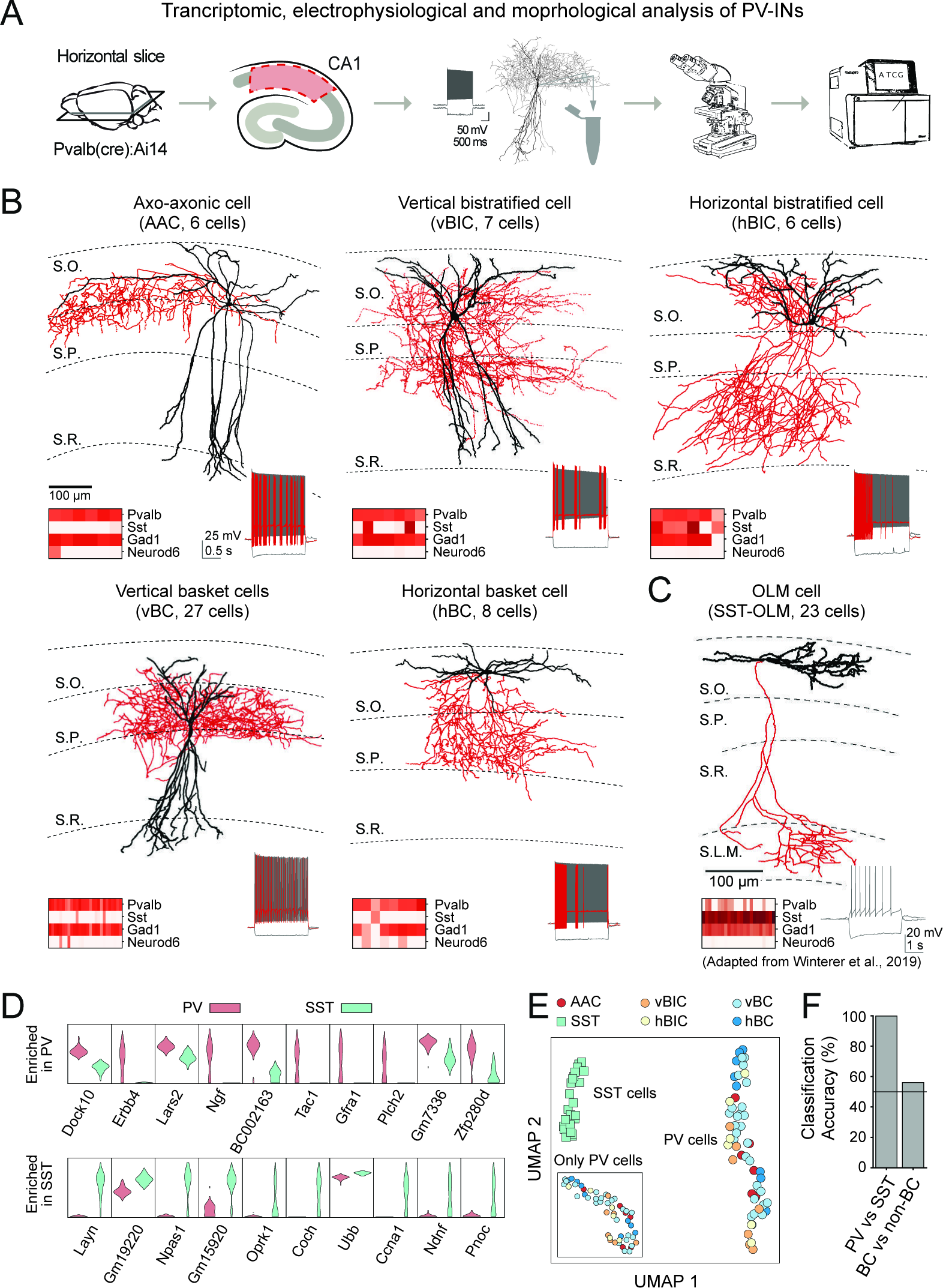
Morpho-transcriptomic profiling of PV-INs. **(A)** Experimental pipeline: tdTomato+ PV-INs were recorded in acute brain slices. Their mRNA was processed for sequencing once the cell’s morphology was confirmed. **(B)** Example morphological reconstructions show each of the five morphological PV types. Insets show electrophysiological responses for hyperpolarizing and depolarizing current injection pulses, and expression of marker genes in single cells (in columns). **(C)** Example morphological reconstruction of an SST-OLM type cell adopted from Winterer et al. (2019). **(D)** Violin plots show expression of top 10 genes that are enriched in PV versus SST-OLM (upper plots) and in SST-OLM versus PV cells (lower plots). **(E)** Plot shows transcriptomic data-based dimension reduction using UMAP. Symbols correspond to different PV and the SST-OLM types. Insert shows UMAP for only PV-INs. **(F)** Bar plot shows classification accuracy for random-forest-based PV versus SST-OLM and for BC versus non-BC discriminations.

To investigate the relation between transcriptomic and morphological cell identities, we collected single-cell RNA-seq data from morphologically identified PV-INs in the CA1 and tested the following three hypotheses.

First, consistent with the view of morphological continuity, a seminal single-cell RNA-seq study on hippocampal CA1 interneurons found that transcriptomically, PV-INs comprise a largely continuous population, divided into two transcriptomic types of approximately equal prevalence, *Pvalb.Tac1* (268 cells) and *Pvalb.C1ql1* (211 cells; Harris et al., 2018; GSE99888; hereafter we refer to this as the ‘CA1-IN study’). In absence of morphological information, this study suggested that these populations represent the AAC and combined BC/BIC types, respectively. This introduces the hypothesis that transcriptomic profiles directly correspond to morphological cell types in addition to other modalities, such as biophysical features.

Second, a central hypothesis suggests that synaptic CAMs determine neuronal connectivity and thus neuronal identity (Sperry, 1963; de Wit and Ghosh, 2016; Südhof, 2017). Consistent with this hypothesis, single-cell transcriptomics have revealed specific CAM expression in multiple neuronal types (e.g. Földy et al., 2016; Shekhar et al., 2016; Paul et al., 2017; Tasic et al., 2018; Mayer et al., 2018; Zheng et al., 2019). Furthermore, our research revealed significant CAM differences between cell types of different developmental origins (Lukacsovich et al., 2019), including the MGE-derived PV and CGE-derived CCK interneurons in CA1 (Földy et al., 2016). However, these studies have also highlighted a higher than expected CAM homogeneity among cell types with the same developmental origin. This has led to the question of whether CAM diversity within a single neuronal family, such as PV-INs, could account for distinct connectivity among different subtypes.

Third, transcriptomic surveys have proposed that transient transcriptomic cell states exist, which may represent a cell’s progress through a developmental trajectory (Le Manno et al., 2018; Mayer et al., 2018; Mi et al., 2018) or a consequence of neuronal activity (Tasic et al., 2018). In the hippocampus, changes during circuit maturation, the development of GABAergic inhibition shows dynamic (Banks et al., 2002; Yu et al., 2006; Fazzari et al., 2010; Salesse et al., 2011) and parvalbumin protein levels continue to increase (Wu et al., 2014). In PV-INs specifically, Er81 (Dehorter et al., 2015) and ErbB4 (Dominguez et al., 2019) levels were found to correlate with activity and plasticity. In fast-spiking interneurons in the cortex, extensive synaptic regulations have been described to take place in the first 4 postnatal weeks (Luhmann and Prince, 1991), which are correlated with transcriptional regulation of thousands of genes (Okaty et al., 2009). These findings give rise to the hypothesis that cell type-specific transcriptional changes regulate the postnatal development of hippocampal PV-INs.

Using morphology-based gene selection, our results help to clarify the transcriptomic identity of morphological PV types. Additionally, they confirm homogenous CAM expression across the whole PV population, which is independent of the cells’ axonal projection or morphological identity. Finally, transcriptomic changes in PV-INs were assessed over the course of circuit maturation, which reveals a surprisingly sharp transcriptomic transition 3 weeks after birth that includes a rapid and stable onset of hemoglobin gene expression.

## Results

### Morpho-transcriptomic characterization of PV-INs

To generate electrophysiological, morphological, and transcriptional data from PV-INs, we performed patch-clamp recordings on cells in the CA1 region of the hippocampus, in brain slices prepared from PV-Cre::Ai14 mice. During recordings, cells were stained with biocytin, which allowed for post-hoc morphological analysis, and after patch-clamp recordings the cytosol was aspirated for subsequent RNA-seq (**Fig. 1A**). Only cells that could be morphologically characterized as stereotypical AAC, vBC, hBC, vBiC or hBiC would be further processed for sequencing (Földy et al., 2016; **Fig. 1B**). From mice that were at least 21 days old, we recorded 195 cells, of which 54 cells were classified as either of the five PV types and passed bioinformatic quality control (6 AAC, 7 vBIC, 6 hBIC, 27 vBC and 8 hBC; see **Methods**). To complement our data, we included our SST-OLM data set adapted from Winterer et al. (2019) as a control MGE-derived, but non-PV hippocampal interneuron type (**Fig. 1C**).

Transcriptomic analysis showed consistent expression of GABAergic marker *Gad1* and absence of glutamatergic marker *Neurod6* in both PV and SST-OLM cells and expression profiles of specific interneuron markers *Pvalb* and *Sst* were consistent with cell type identity (**Fig. 1B** and **C**). Using *edgeR* (McCarthy et al., 2012), 124 genes were found to be enriched in PV compared to SST cells, and 47 genes were higher expressed in SST compared to PV-INs with *p*-adjusted (from here on referred to as *p*) <0.05 and fold change >2 (top 10 genes of each comparison are shown in **Fig. 1D**; see also **Fig. S1**). Consistent with previous studies, *Erbb4* and *Tac1* were enriched in PV and *Pnoc* in SST cells (Neddens and Buonanno, 2010; Harris et al., 2018; Mayer et al., 2018; Tasic et al., 2018). Using UMAP (Uniform Manifold Approximation and Projection; McInnes et al., 2018), we performed dimension reduction on the transcriptomic data, which revealed that PV and SST cells separated into different groups, but morphological PV types did not distinctly cluster, even when plotted without SST cells (**Fig. 1E**). In addition, random forest classification accurately classified cells as PV or SST type (99.8% accuracy) but could not further distinguish morphological PV types (56.2% accuracy in case of the BC versus non-BC comparison, where sample numbers were sufficient to define separate training and testing sets; **Fig. 1F**).

To further analyze any transcriptional differences in morphological PV types, we applied proMMT (Probabilistic Mixture Modeling for Transcriptomics; a combined workflow for gene selection-based transcriptomic clustering; Harris et al., 2018) that was previously used to analyze CA1-IN data. ProMMT yielded four transcriptional clusters, which we visualized using nbt-SNE (negative binomial t-SNE; Harris et al., 2018; **Fig. 2A**). Notably, these four transcriptional groups could not be split into distinct clusters in two-dimensional space, even when other dimension reduction techniques were applied (PCA, t-SNE, Flt-SNE and UMAP; **Fig. S2**), nor did they correspond with the morphological types. To assess the transcriptional signatures of PV-INs in a larger context, we included the above referenced CA1-IN data. Meta-analysis of this data revealed that our cells showed unconstrained mapping onto the two populations that were identified as PV cells in the original study. Using six different gene selection methods, our PV-INs could be consistently mapped onto either of the two PV associated continents (**Fig. 2B** and **C**). However, morphological types did not correlate with the *Pvalb.Tac1* (presumed AAC) or *Pvalb.C1ql1* types (presumed BC/BIC population; **Fig. 2C**), suggesting that the clustering in the CA1-IN study did not arise from the 3 main PV types. In conclusion, these findings further indicate that morphological PV types are not majorly distinct transcriptionally from one another.

**Fig. 2.**
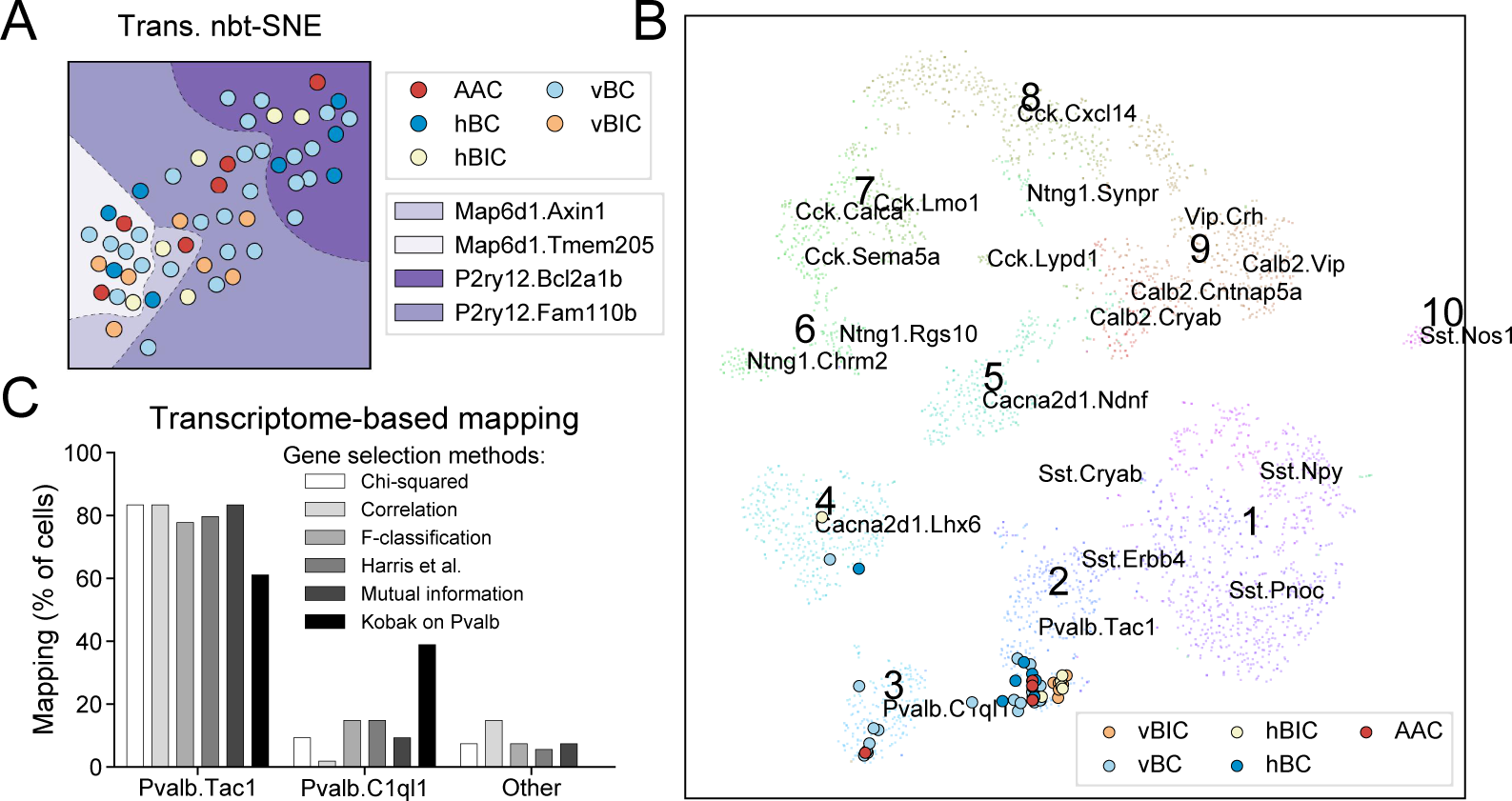
Transcriptomic properties of PV-INs. (A) Plot shows unsupervised, transcriptomic data-based dimension reduction using nbt-SNE (Harris et al., 2018). Single cells are labeled with circles, which are colored according to morphological classification. Dashed lines show transcriptomic classification using proMMT (Harris et al., 2018). **(B)** Plot shows mapping of PV-INs of this study onto the CA1-IN data set (Harris et al., 2018). Cell type labels, e.g. *Pvalb.Tac1*, are also imported from the original CA1-IN study. Gene selection for mapping was performed using the method described in Kobak et al. (2019). **(C)** Quantification of mapping efficacy using six different gene selection methods (see Methods for details)

### PV-INs comprise a biophysically homogenous population

To assess whether yet un-covered biophysical differences further characterized morphological PV types, we quantified 10 electrophysiological parameters, including passive (e.g. input resistance and membrane capacitance) and active (e.g. properties of single and train AP firing) membrane properties. However, pair-wise comparisons for each electrophysiological property between the 5 morphological PV types (total of 100 comparison) did not reveal statistically significant differences, with the only exceptions of membrane capacitance between hBIC and vBC types (*p* = 0.015, two-sided Welch’s t-test) and input resistance between hBC and vBC types (*p*=0.012; **Fig. 3A**). To further corroborate, we used UMAP for dimension reduction to assess potential clustering among the electrophysiological parameters. Such clusters may arise due to a combination of electrophysiological differences, which are not necessarily significant, but together differentiate morphological or transcriptomic types. We considered the following biologically relevant scenarios: 2 ‘dendro-morphological’ types (vertical versus horizontal distinction), 3 ‘axo-morphological’ types (AAC, BC, and BIC), 4 proMMT transcriptomic types (as in **Fig. 2A**) and the 5 morphological PV types. However, cells did not cluster along any of these distinctions (**Fig. 3B**), and clustering between any two of the electrophysiological parameters were also lacking (**Fig. S3**). To conclude, these results show a pronounced biophysical homogeneity among PV-INs, regardless of their morphological or transcriptional differences.

**Fig. 3.**
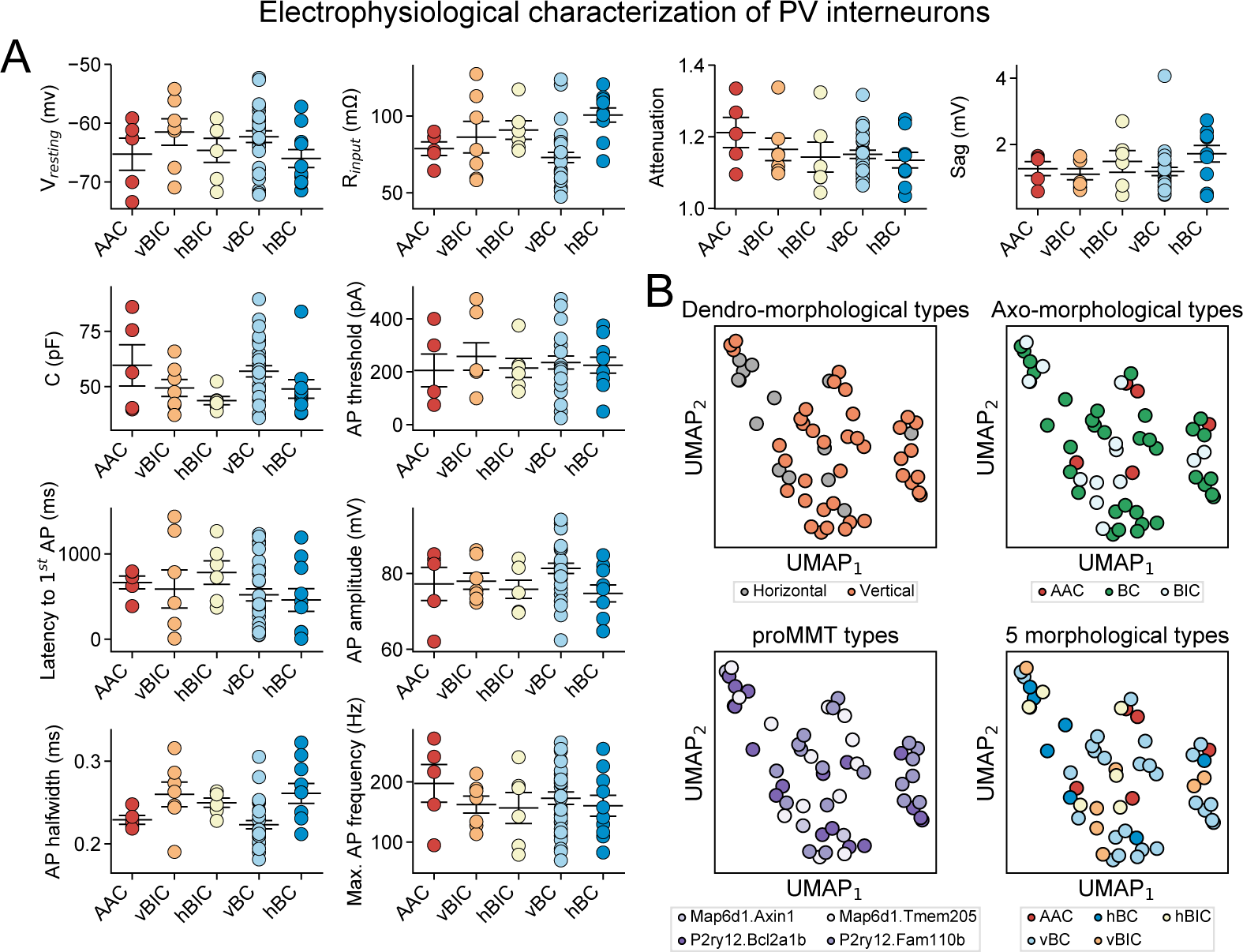
Electrophysiological properties of PV-INs. (A) Plots show electrophysiological properties measured in the 5 morphological PV types. Circles represent single cells. **(B)** All panels show dimension reduction on electrophysiological data using UMAP. Circles represent single cells and are colored to show the 2 dendro-morphological, 3 axo-morphological, 4 proMMT transcriptomic, and 5 morphological PV types.

### Transcriptomic definition of morphological PV types

To transcriptomically define morphological types, we examined gene expression differences among the 5 morphological types, as well as among the axo- and dendro-morphological types. The 7 differentially expressed genes among the 5 morphological types (criteria were at least 2-fold difference in expression level and *p* < 0.05 between any two types using quasi likelihood test) included *Akr1c18*, *Kcng4*, *Synpr* (all three in AAC versus vBIC comparison; *Synpr* and *Akr1c18* being enriched in vBIC, whereas *Kcng4* in the AAC type), *Esyt1* (in hBC versus vBC), *Npy* (in vBIC versus hBC and vBC), and *Sst* (hBIC versus vBC; **Fig. 4A**; see Discussion for details on these genes). PCA on these genes revealed a graded distribution of morphological PV types and most notably the separation of BIC type (**Fig. 4A**, lower panel). Comparison of axo-morphological types revealed already detected genes that distinguished BIC-s by enrichment (i.e. *Sst*, *Npy*, *Synpr*, *Akr1c18*), but also novel genes (*Gpc6*, *Cemip*, *Srgn*, *Pthlh*, and *Trpc3)* specifically lacking from BIC types (**Fig. 4B**). Finally, the comparison of dendro-morphological types revealed, among others, enrichment of *Ndufa1*, *Slc37a3* and *Tuft1* (*p*=0.011, 0.021 and 0.047) in vertical and *Zfx*, *Slc7a7*, and *Ctso* (*p*=0.046, 0.017, and 0.046) in horizontal types (**Fig. 4C**; genes with *p*<0.05 are shown). Using an extended set of 88 genes from all three morphology-based comparisons (*p*<0.15; see SI for complete list), PCA now showed clear separation of the morphological BIC and AAC/BC types (**Fig. 4D**). Moreover, it highlighted a distinction between the hBIC and vBIC, and although to a lesser degree, between the vBC and hBC types (for implementation of support vector machine classification on this same problem and its conclusions, see **Fig. S4**). In conclusion, despite the transcriptomic homogeneity of PV-INs, morphology-based transcriptomic analyses revealed gene-expression differences that separately identify the vBIC and hBIC types but did not further differentiate AAC and BC types.

**Fig. 4.**
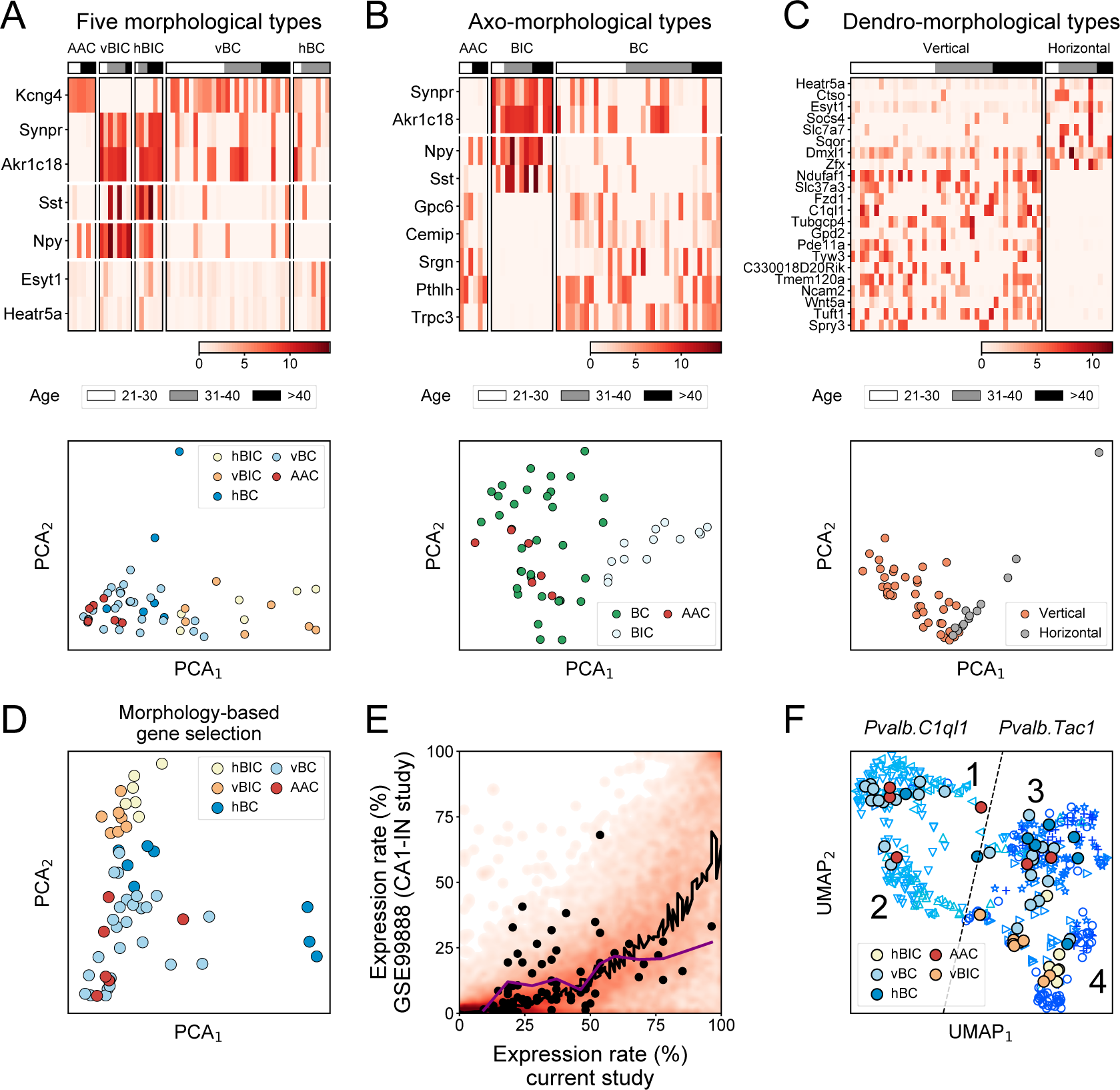
Transcriptomic definition of morphological PV types. **(A-C)** Heat maps (upper plots) of differentially expressed genes (using *edgeR*, fold difference>2, *p*<0.05) between the 5 morphological PV types (panel A), 3 axo-morphological types (panel B), and 3 dendro-morphological types (panel C). Cells are ordered according to age. PCA plots (lower plots) were made using the differentially expressed genes. **(D)** PCA plot of cells, colored by the 5 morphological PV types, using an extended set (with a cutoff of *p*<0.15) of n=88 differentially expressed genes from A-C. **(E)** Comparison of expression rate of genes in PV-INs from this current versus the CA1-IN study, displayed as a heat map. Black line marks loess regression fit. Black points label the 88 differentially expressed genes (with a cutoff of *p*<0.15) from A-C. **(F)** UMAP based embedding of PV-INs from the CA1-IN study (Harris et al., 2018), and mapping the PV-INs of this current study onto the UMAP embedding using the 88 differentially expressed genes (with a cutoff of *p*<0.15) for A-C. Symbols refer to the following transcriptomic subtypes, as described in the original study: ▷ *Pvalb.Tac1.Akr1c18*, ◁ *Pvalb.Clql1.Cpne5,* △ *Pvalb.C1ql1.Npy,*▽*Pvalb.Clql1.Pvalb,* + *Pvalb.Tac1.Nr4a2,* ○ *Pvalb.Tac1.Sst,* and ⋆ *Pvalb.Tac1.Syt2*.

Next, we evaluated the expression of the 88 morphology-associated genes in the CA1-IN data set. We found that the majority of genes, including these 88, were enriched in our data (**Fig. 4E**) and several of them were not or very rarely detectable in the CA1-IN data set (7 genes were not detected, another 25 were detected in at most 20 out of 479 cells, and yet another 23 were detected in less than 10% of CA1-IN cells; **Fig. S5**). This discrepancy, which may be due to the more than a 100-fold difference in alignment depths (10 million versus 0.1 million reads per cell in this and in the CA1-IN study), suggested a sub-optimal recovery of morphologically relevant genes in the CA1-IN study. Even with this caveat, UMAP dimension reduction using the 88 genes still introduced finer distinctions within the CA1-IN *Pvalb.Tac1* and *Pvalb.C1ql1* types while preserving their global composition (**Fig. 4F**). Subsequent mapping of our PV cells onto this refined CA1-IN transcriptomic map lead to the following observations: 1) *Pvalb.C1ql1* islands 1 and 2 as well as the bridge between island 2 and 3 were mapped by vertical cells, 2) *Pvalb.Tac1* island 3 was mapped by both BC (vertical and horizontal) and AAC, but not BIC, type cells, and 3) *Pvalb.Tac1* island 4 was mapped by only BIC (vertical and horizontal) type cells (**Fig. 4F**). To sum, our results revealed sub-optimal conditions to resolve all morphological PV types in this transcriptomic map. Nevertheless, they indicate that *Pvalb.C1ql1* represented cells with vertical dendrites, *Pvalb.Tac1* represented a mixed pool of vertical and horizontal type cells, and that the *Pvalb.Tac1.Sst* and *Pvalb.Tac1.Akr1c18* subregions (island 4) appeared to consist of exclusively BIC type cells.

### CAM homogeneity among morphological PV types

Although our results already revealed a pronounced transcriptomic homogeneity among morphological PV types, we further investigated the expression of CAMs due to their importance in specifying neuronal connectivity and their close relation to neuronal identity. However, as predicted by the above analyses, CAM expression (based on 405 genes; see Földy et al., 2016) was highly uniform among the 5 morphological PV types (**Fig. 5A**). Although pairwise comparisons (using *edgeR*, fold difference>2, *p*<0.05) revealed fine distinctions between morphological PV types, such as enrichment of *Icam1* in hBC and lack of *Ptprt* in vBC type (**Fig. 5B**, upper panel), and between pooled PV cells and SST cells (**Fig. 5B**, lower panel), CAM based similarity comparisons showed uniformity among the different PV types (**Fig. 5C**). Meanwhile, PV-INs consistently lacked expression of CAMs that were previously associated with CA1 regular spiking (RS; presumed CCK population) and CA1 pyramidal types (PYR; Földy et al., 2016; **Fig. 5D**), corroborating specific CAM expression when compared to developmentally different neuronal families. To sum, we conclude that CAM expression is highly uniform across the entire PV-IN population.

**Fig. 5.**
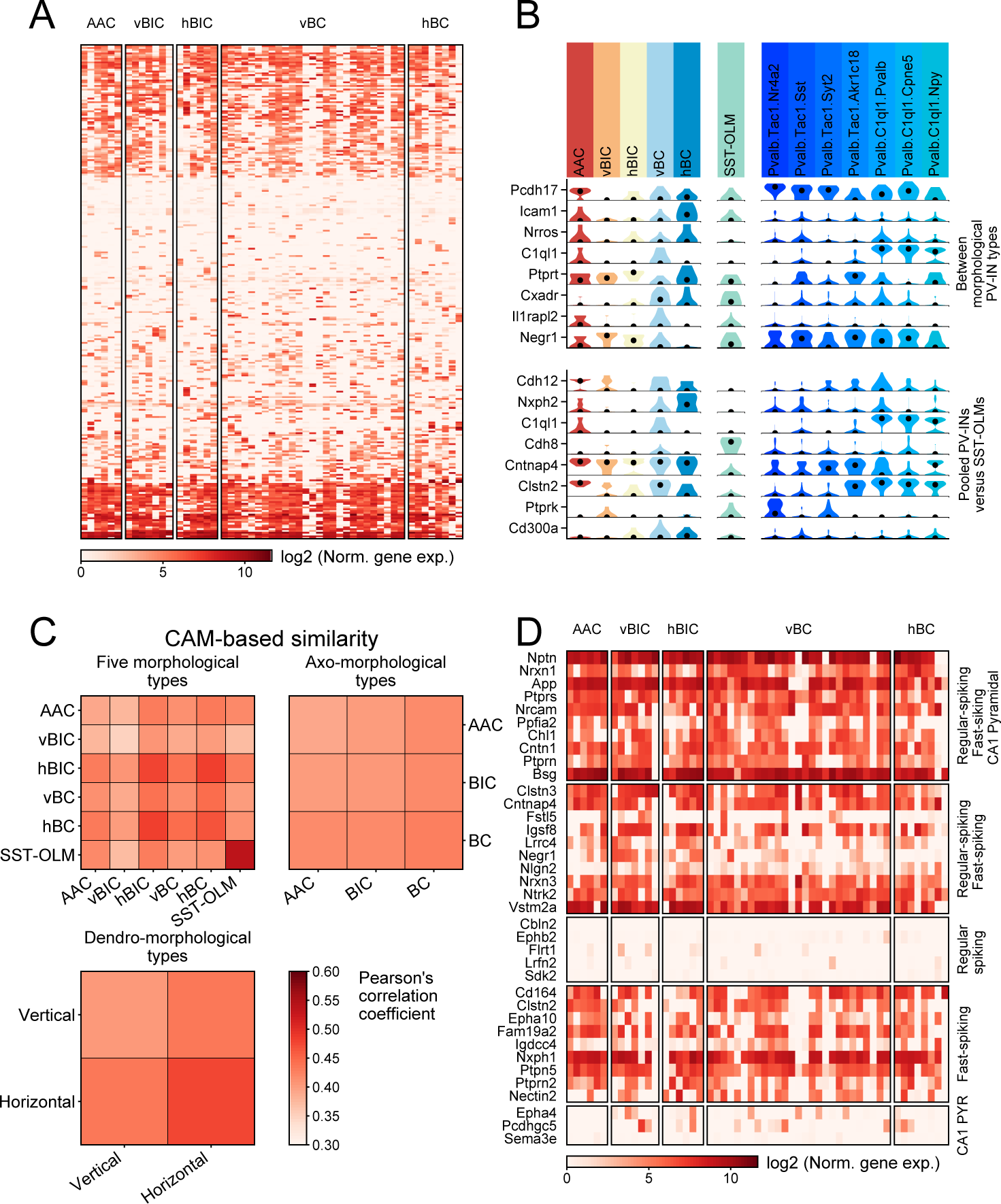
Analysis of CAM expression in morphological PV types. (A) Heat map showing all CAMs expressed in at least 3 PV-INs, in PV (grouped by morphological type) and SST-OLM cells. Genes are ordered via
hierarchical clustering based on their expression in PV-INs. (B) Violin plots show the top 8 most significantly differentially expressed CAMs (using *edgeR*, fold difference>2), between morphological PV types (top panel), and between pooled PV-INs and SST-OLM cells (bottom panel). (C) Similarity matrices between the 5
morphological (top left), 3 axo-morphological (top right), and 2 dendro-morphological PV types (bottom left).
Similarity scores were measured based on only CAM expression and using average of Pearson’s correlation
coefficients of cells. (D) Heat map shows expression of genes that were previously described to be (1) ubiquitous
in CA1 pyramidal (PYR), fast-spiking (FS; presumed PV-INs), and regular-spiking cells (RS; presumed CCKINs);
or specifically expressed in (2) FS and RS cells; (3) RS cells; (4) FS cells; or in (5) PYR cells (Földy et
al., 2016).

### Functional maturation of PV-INs

To extend our analysis into an earlier developmental domain, such as the critical period of interneuron plasticity (Banks et al., 2002; Salesse et al., 2011; Callahan et al., 2013; Domínguez et al., 2019), we collected additional cells from younger than 21 days old (<P21) animals. To make this analysis more focused, we emphasized collecting data from the vBC type, by patching tdTomato+ cells within the pyramidal cell layer, which consisted the majority of our >P21 data set (in mice, PV and thus tdTomato+ expression first appears at ∼P10, which limits cell collection from earlier time points using this approach). In this manner, we complemented our existing data set with an additional n=19 vBC and analyzed a combined number of 46 vBC type PV-INs collected between P10 and P77.

Morphological analysis showed that <P21 vBC type PV-INs already display fully developed axonal and morphologic features (**Fig. 6A**). Sholl analysis on dendritic arborizations showed similar patterns between <P21 and >P21 cells (**Fig. 6B**), and dendritic and axonal lengths were not significantly different (*p*=0.07 and 0.8, respectively; **Fig. 6C** and **D**), where P21 was used as an arbitrary cut-off. However, in accordance with an on-going functional maturation of the circuit, electrophysiological properties displayed age-dependent changes. Specifically, we found shorter AP half-width (*p*=0.0013), lower average AP firing frequency (*p*=0.013), and faster AP firing attenuation (*p*=3.2×10^-6^, two-sided Mann-Whitney test was used for these comparisons; **Fig. 6E**) in <P21 versus >P21 cells. While correlated with age, these physiological differences could not separately cluster differently aged PV-INs (**Fig. 6F**). To elaborate in the transcriptomic domain, we examined ion channel-coding genes, and identified only one significant change, the decreased expression of the potassium channel subunit *Kcnq3* in P>21 (**Fig. S6**). We hypothesize, but did not test further, that this subunit which underlies the M current (Wang et al., 1998) contributed to the observed physiological changes.

**Fig. 6.**
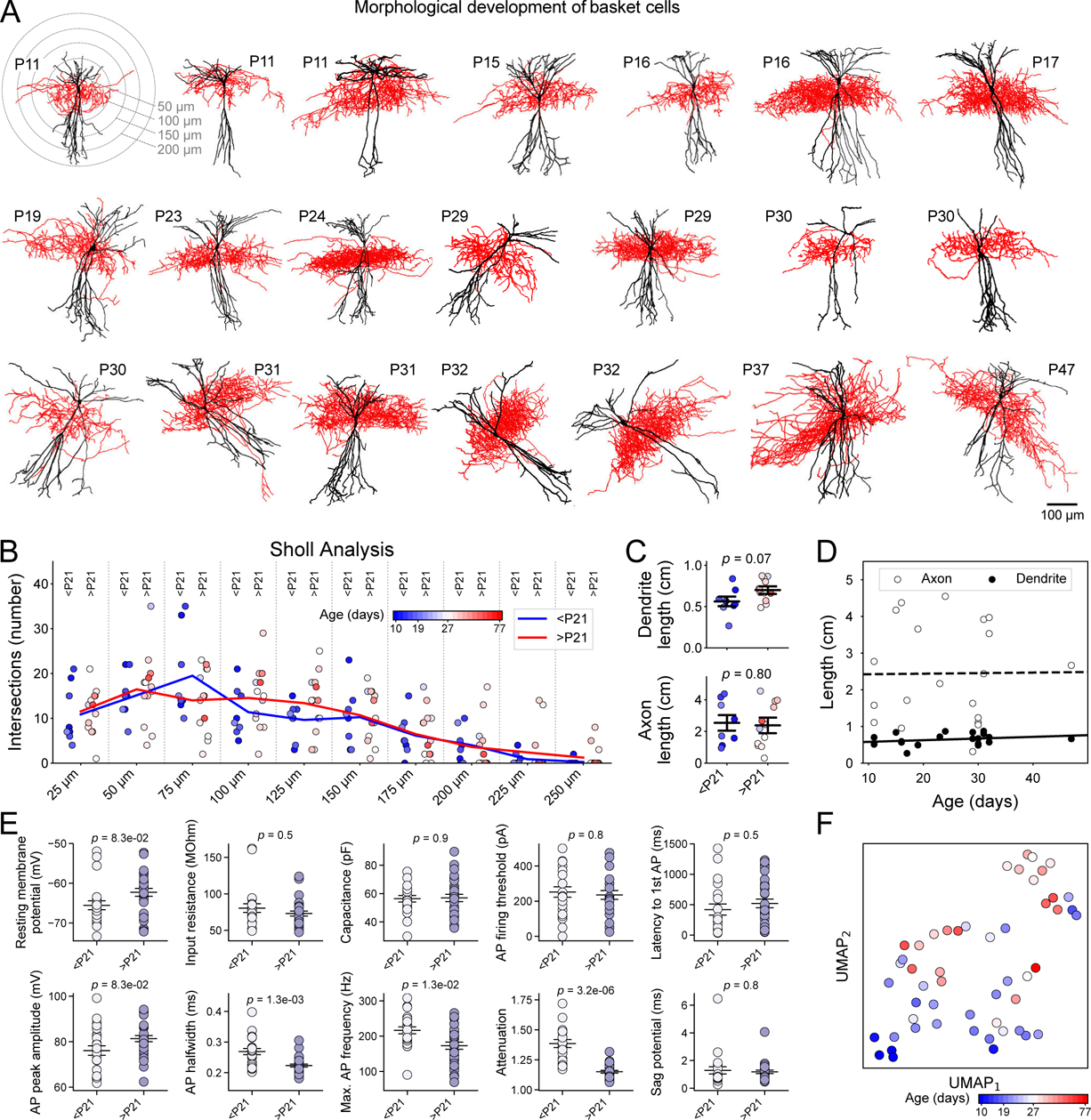
Morphological and electrophysiological analysis of vBC type PV-INs during circuit maturation. (A) Morphological reconstructions show vBC type PV-INs at different ages. First image shows Sholl analysis
for dendritic branching based on concentric 3D spheres (plot shows circles for clarity). (B) Plot shows the
number of intersections at different distances from the soma of n=8 <P21 and n=13 >P21 vBC. Circles denote individual cells. None of the statistical comparisons (Welch’s t-test) revealed a p value smaller than 0.125. (C) Plots show the total dendritic (left) and axonal (right) length of <P21 and >P21 vBC type cells. P values are shown on top and were determined using Welch’s t-test. (D) Scatter plots and linear regression fits of axon and dendrite length against cell age. Neither value was shown to correlate with age (lowest p value and highest rsquared for fit were 0.241 and 0.075 respectively). (E) Plots show electrophysiological properties of vBC type cells in n=23 <P21 versus n=29 >P21 mice, together with corresponding p value (Welch’s t-test). Attenuation (p=3.2×10-6), AP halfwidth (p=0.0013) and maximum AP frequency (p=0.013) show significant changes. (F) UMAP based dimension reduction of electrophysiological properties of vBC type cells. All cells represent the vBC type and are colored by age.

### Rapid transcriptomic changes in PV-INs between P21 and P25

Using two independent bioinformatic approaches, we then examined whether transcriptomic changes other than ion channels corresponded to PV cell maturation. First, we used a sliding-window approach, only considering whether a gene was expressed or not, independent of its expression level (**Fig. 7A** and Methods). Second, we used Monocle (Qiu et al., 2017), which relies on expression levels and calculates the cells’ pseudo time best correlating with age and finds genes whose expression level significantly correlate with this (**Fig. S7**). As a result, we found a surprisingly short period between P21 and P25, which was marked by pronounced down-regulation of n=48 genes (Monte Carlo on Gini impurity, *p*<0.1; Methods; see supplementary excel file for complete list) and up-regulation of n=5 genes (Monte Carlo on Gini impurity, *p*<0.1; **Figs. 7A**). This pattern was robust and separately clustered cells by their age (**Fig. 7B**) and moreover, using random forest classification, predicted whether the animals’ age was below P21 or above P25 (**Fig. 7C**). Additional gene ontology (GO) analysis revealed that while the down-regulated genes included multiple different families, none of the ontologies changed with a *p* value lower than 0.19 (**Fig. 7D**).

**Fig. 7.**
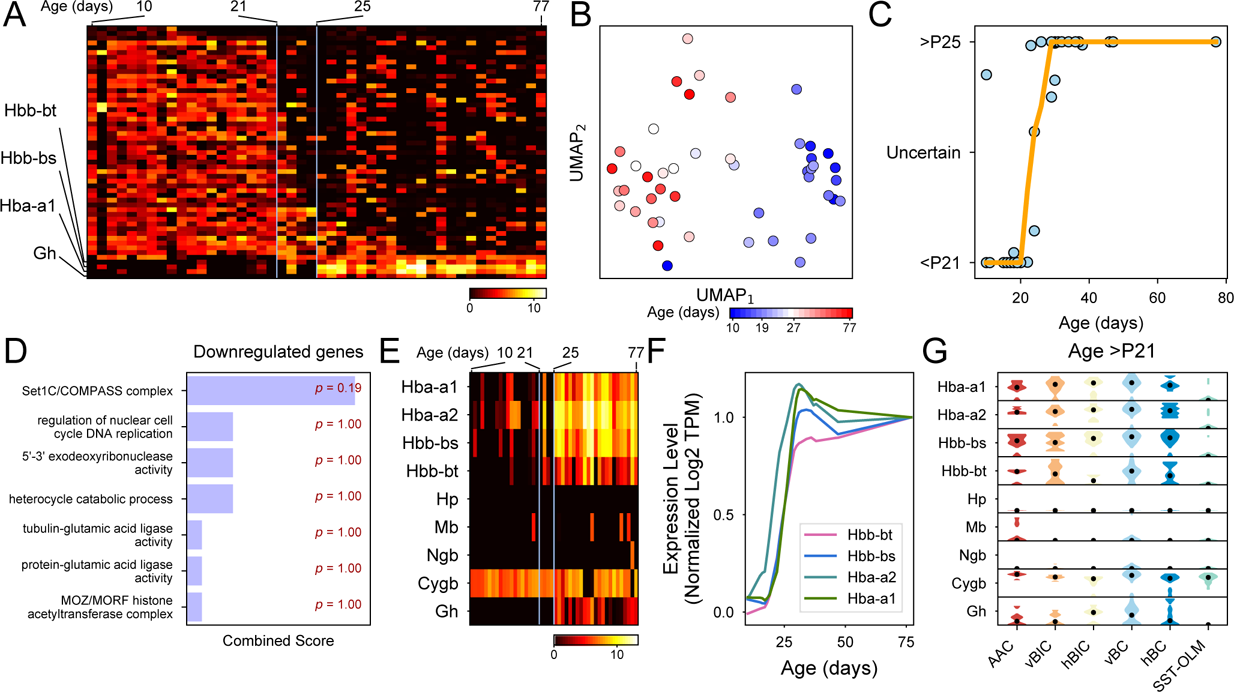
Age-dependent transcriptomic changes and onset of hemoglobin expression in PV-INs. (A)
Expression levels of genes showing a statistically significant (p<0.10) transition using Gini impurity (see
Methods). Genes (rows) ordered by the day of transition, cells (columns) are ordered according to age. (B)
UMAP based on genes from panel A. All cells represent the vBC type and are colored by age. (C) Plot of
consistency with which a cell’s age can be predicted using Random Forest Classifier based on genes from panel
A (see Methods), fitted with a loess curve. (D) Plot shows top GO scores using enrichR on downregulated genes
from panel C. (E) Heat map shows the expression of hemoglobin and related genes in vBC type cells (columns),
which are sorted by age left to right. (F) Normalized loess fits of hemoglobin expression levels versus cell age
in vBC type cells. (G) Violin plots show expression of haemoglobin and related genes from panel E only
considering >P21 PV-INs and SST-OLM cells.

### Unexpected onset of hemoglobin expression in PV-INs

Among the up-regulated genes we found *Gh* (or growth hormone 1), and surprisingly *Hba-a1*, *Hbb-bt* and *Hbb-bs*, which are all hemoglobin (Hb) subunit-coding genes, and additionally one pseudogene (*GM443889*). Although not detected by our initial analysis, we also examined the expression of *Hbb-a2* (another key component of functional Hb tetramers) and that of *Mg* (myoglobin), *Ngb* (neuroglobin) and *Cytb* (cytoglobin), which all display oxygen binding properties similar to Hb (Ascenzi et al., 2014). *Hbb-a2* followed a similar expression pattern as the other Hb-coding genes. By contrast, *Mg* and *Ngb* were not expressed, whereas *Cytb* was highly expressed, without regard to age (**Fig. 7E, F** and **S7**). We also explored the expression of genes which are known to regulate Hb expression. We found a lack of canonical Hb transcriptional regulators GATA-1 and −2 (Katsumura et al., 2013; Ascenzi et al., 2014), but stable (age independent) expression of *Hif1a*, a hypoxia response-related transcriptional factor that also controls Hb expression (**Fig. S7**). Finally, this unexpected onset of Hb expression was characteristic to all PV, but not SST, types (**Fig. 7G**). This specificity is also supported by ISH staining of Hb subunits from the Allen Mouse Brain Atlas (**Fig. S7**).

## Discussion

Multiple studies already suggest that physiological features can be inferred from single-cell transcriptomic data (Okaty et al., 2009; Cadwell et al., 2016; Földy et al., 2016; Fuzik et al., 2016; Muñoz-Manchado et al., 2018; Luo et al., 2019; Oláh et al., 2019; Winterer et al., 2019; Zheng et al., 2019). By contrast, the ability to infer neuronal morphology from transcriptomic information has not yet been established. Large scale transcriptomic assays have previously described continuously varied gene expression among and within cell types, in which clusters may split (discreteness) or merge (continuous variation), depending on gene detection, cell sampling numbers, and noise estimates or statistical criteria (Tasic et al., 2018). For CA1 interneurons specifically, continuous variation was earlier suggested based on a transcriptomic map that also included two distinct transcriptomic PV-IN clusters, appointed to be presumed AAC and BC/BIC types (Harris et al., 2018). Here, we sequenced mRNA from morphologically identified PV-INs in hippocampal CA1 to relate transcriptomic content to morphology and circuit connectivity.

### Transcriptomic definition of morphological PV types

Using unsupervised feature selection and dimension reduction on the cells’ whole transcriptome, we showed that while PV and SST cells can be transcriptionally separated into their respective populations, known morphological PV types could not be accurately distinguished.

Meta-analysis of the CA1-IN data set with our own revealed that our cells corresponded to the presumed PV population, but did not support the hypothesis that morphological AAC and BC/BIC types defined major transcriptomic types (**Figs. 1-3**). By contrast, supervised gene selection based on morphology confirmed known and revealed novel subtype specific gene expression patterns (**Fig. 4**).

As known markers, we confirmed that *Sst* and *Npy* were enriched in the BIC type (see Katona et al., 2014). As a finer distinction, our data also revealed a complementary expression of the two genes within the BIC population: *Npy* was enriched in vBIC, whereas *Sst* was enriched in hBIC. As novel markers, *Synpr* and *Akr1c18* were enriched in the BIC type, whereas among others *Kcng4*, *Phtlh* and *Trpc3* were enriched in the AAC and BC types. Of these, (1) the correlation of *Akr1c18* with fast- and delay-spiking markers within the *Pvalb.Tac1* population has been highlighted, but without association to morphology (Harris et al., 2018). (2) *Kcng4*, which encodes a modulatory subunit for the potassium channel Kv2.1 (Sano et al., 2002), may represent a highly selective marker for both AAC and BC types. According to the Allen Mouse Brain Atlas, *Kcng4* is present only in a handful of cells in the pyramidal layer of CA1, plausibly suggesting restricted expression in PV-INs. Added to this, our data shows specificity to AAC and BC within PV-INs. (3) *Pthlh* expression was previously found to correlate with fast-spiking property of PV-INs in the dorsal striatum (Muñoz-Manchado et al., 2018). By contrast, our data in CA1 show correlation of *Pthlh* with morphological features. Since the striatum study did not include morphological characterization, it is possible that fast-spiking property in striatal PV-INs also correlated with morphology. (4) Finally, *Trpc3* expression in the BC type may identify a yet unknown mediator of the CCK-induced transient receptor potential (TRP) currents, which we previously measured in BC, but not in BIC, type PV-INs (Lee et al., 2011). Although our data revealed multiple genes that differentiated BIC from AAC and BC types, it did not disclose genes differentiating the AAC and BC types. This contrasts with a previous observation made in the CA3 area, where SATB1 was specifically expressed in the AAC, but not BC, type (Viney et al., 2013).

Our results shed new light onto transcriptomic differences among dendro-morphological types. We identified 14 and 8 genes, which were selectively enriched in vertical and horizontal types, without regard to the cells’ axo-morphological features. Our observations furthermore suggest that dendro-morphological features also need to be considered when interpreting transcriptomic types. Meta-analysis of the CA1-IN data set (Harris et al., 2018) with our own revealed that *Pvalb.C1ql1* likely represented cells with vertical dendrites (we detected *C1ql1* only in vertical, but not in hortizontal, cells, **Fig. 4**), whereas *Pvalb.Tac1* represented a mixed pool of cells with vertical and horizontal dendritic cells. However, we also found that a number of morphological marker genes we detected in our cells were not at all or only infrequently detected in the CA1-IN data. The reason for this discrepancy is unknown, nonetheless limited further parsing of this data set into morphological types. In conclusion, our analyses showed that morphological PV types display a high homogeneity at the whole transcriptome level, but also that specific expression of a handful of marker genes can differentiate the BIC and AAC/BC types from one other.

### Molecular architecture of circuit connectivity

Transcriptomic data collected from morphological PV types allowed us to further test a key theory of brain connectivity which states that synaptic CAMs specify neural connections. While ample evidence supports this hypothesis (de Wit and Ghosh, 2016; Südhof 2018, for reviews), in seeming contradiction, our data revealed a striking homogeneity of CAM expression among the morphological PV types (**Fig. 5**). However, this outcome was not completely unexpected. Transcriptomic studies have revealed major CAM differences between different neuron families, such as between excitatory versus inhibitory cells or between inhibitory cells with different developmental origin (Földy et al., 2016; Tasic et al., 2018; Lukacsovich et al., 2019; Zheng et al., 2019), but CAM diversity appeared to be less pronounced when only cells within individual neuron families, such as PV-INs, were considered (Földy et al., 2016; Lukacsovich et al., 2019). However, supporting evidence demonstrating the cells distinct morphology or connectivity was lacking from these studies. Our current study provides evidence for CAM homogeneity among PV-INs, which exists without regard to their morphological features. As a corollary of this finding, our ability to use mRNA expression-based transcriptomic information to infer circuit connectivity remains limited. It is nevertheless possible that isoform level, non-mRNA, or translational information will prove be sufficient for making such predictions (Que et al., 2019). Alternatively, the input/output connectivity of these cells may be specified exclusively during earlier development, after which key factors become downregulated (Favuzzi et al., 2019), and connectivity patterns are not actively reinforced later in life.

### Switch of transcriptomic states and rapid onset of Hb expression

During the second postnatal week of cortical development, fast-spiking interneurons display intense transcriptomic changes that involve thousands of genes (Okaty et al., 2009). Ion channel coding genes change their expression with up to 10-100-fold magnitude, which coincide with profound electrophysiological maturation of cells. By contrast, our analysis did not register electrophysiological, transcriptomic or morphological changes at a similar scale, suggesting that, in hippocampus, intrinsic maturation of PV-INs is largely completed by P10. Our data, however, revealed another wave of transcriptomic regulation, which occurred at a later time window (between P21-P25) and was restricted to a smaller number (∼50) of genes (**Fig. 7**). Most of the genes displayed downregulation and represented functionally diverse families. Importantly, these did not include CAMs, suggesting that any potential downregulation within this gene family occurred during earlier development, in accordance with a seeming morphological completion after P10 (**Fig. 6**).

By contrast, fewer genes were upregulated, most of which encoded Hb subunits. While this was unexpected, Hb expression has been previously demonstrated in a limited number of neuronal types, including A9 dopaminergic (Biagioli et al., 2009) and unidentified type of cortical, hippocampal and cerebellar cells (Wu et al., 2004; Schelshorn et al., 2009; Richter et al., 2009). The onset of Hb expression characterized all morphological PV types. The role of Hb expression in neurons remains controversial (Biagioli et al., 2009; Schelshorn et al., 2009; Richter et al., 2009). In hypoxia, Hb appeared to be neuroprotectant by rendering cells into an oxygen privileged state (Schelshorn et al., 2009). However, neurodegenerative effects of Hb expression were also proposed (in aging, Blalock et al., 2003; by promoting Aβ oligomerization, Wu et al., 2004; by toxic Hb aggregate formation, Richter et al., 2009; and by learning impairments, Codrich et al., 2017). In hippocampus specifically, chronic stress lead to significant downregulation of Hb genes (Andrus et al., 2012), whereas early-life iron deficiency anemia altered the development and long-term expression of parvalbumin and perineuronal nets (Callahan et al., 2013). While our results do not clarify the role of Hb gene expression in neurons, they make a novel observation that specifically implicates PV-INs as a cellular substrate behind Hb-associated network effects.

### Summary

This study performed a combined analysis of hippocampal PV-INs in the electrophysiological, morphological and transcriptomic domains. Outcomes identified transcriptomic signatures that discern morphological PV types but corroborated an overall transcriptomic homogeneity among the entire PV population. Furthermore, this study provides evidence for a lack of differentiating CAMs (as defined by mRNA based transcriptomic readout) among differently wired cell types. Finally, results of this study demonstrate a switch of transcriptomic states and rapid onset of Hb expression, which may directly relate to PV-IN pathology behind certain neurodevelopmental and neuropsychiatric disorders (Marin 2012; Wöhr et al., 2015).

## Materials and Methods Animals

All animal protocols and husbandry practices were approved by the Veterinary Office of Zürich Kanton. The University of Zurich animal facilities comply with all appropriate standards (cages, space per animal, temperature, light, humidity, food, water) and all cages were enriched with materials that allow the animals to exert their natural behavior. Both males and females were used for all experiments. Animals were sacrificed from P10 and older. The following lines were used in this study: (1) PV-CRE and (2) Ai14: B6.Cg-Gt(ROSA)26Sor<tm14(CAG-tdTomato)Hze>/J Stock No: 007914.

### Electrophysiology

Hippocampal slices (300 µm thick) were prepared from P10 and older mice, and incubated at 34°C in sucrose-containing artificial cerebrospinal fluid (sucrose-ACSF) (85 mM NaCl, 75 mM sucrose, 2.5 mM KCl, 25 mM glucose, 1.25 mM NaH_2_PO_4_, 4 mM MgCl_2_, 0.5 mM CaCl_2_, and 24 mM NaHCO3) for 0.5 h, and then held at room temperature until recording. Cells were visualized by infrared differential interference contrast optics in an upright microscope (Olympus; BX-51WI) using a Hamamatsu Orca-Flash 4.0 CMOS camera. Recordings were performed using borosilicate glass pipettes with filament (Harvard Apparatus; GC150F-10; o.d., 1.5 mm; i.d., 0.86 mm; 10-cm length) at 33 °C in ACSF (126 mM NaCl, 2.5 mM KCl, 10 mM glucose, 1.25 mM NaH_2_PO_4_, 2 mM MgCl_2_, 2 mM CaCl_2_, and 26 mM NaHCO_3_) with a standard intracellular solution (95 mM K-gluconate, 50 mM KCl, 10 mM Hepes, 4 mM Mg-ATP, 0.5 Na-GTP, 10 mM phosphocreatine; pH 7.2, KOH adjusted, 300 mOsm). All recordings were made using MultiClamp700B amplifier (Molecular Devices), and signals are filtered at 10 kHz (Bessel filter) and digitized (50 kHz) with a Digidata1440A and pClamp10 (Molecular Devices).

### Identification of cell types

Neurons were identified by fluorescent labeling in hippocampal brain slices prepared from Pv-Cre::Ai14 mice. Fluorescence-labeled cells were variably present in all hippocampal strata. During recording, cells were filled with biocytin (Sigma-Aldrich, 2%) for subsequent *post hoc* visualization of axons. After collection of cytosols, brain slices were fixed in 4% Paraformaldehyde (Sigma-Aldrich) overnight and subsequently processed for immunostaining with streptadivin-alexa Fluor 488 conjugate (Invitrogen, Thermo Fisher Scientific). Only those cells were included, where staining revealed axonal and dendritic arborization. Out of 309 cells recorded for the whole study, 182 cells were not included based on insufficient staining of the axons, which could be caused by technical artifacts such as brain slice preparation or staining issues. Furthermore, 37 cells could not be unambiguously classified as either of the five PV types and finally, 15 cells did not pass quality control after single-cell RNA sequencing. Cells are listed by cell name, cell type, age in supplementary excel file.

### Single-cell RNA sequencing

#### * Sample Collection

Methods and practices are identical as we described before in Földy et al. (2016) and Winterer et al. (2019). To minimize interference with subsequent molecular experiments, only a small amount of intracellular solution (∼1 µl; not autoclaved or treated with RNase inhibitor) was used in the glass pipette during electrophysiological recordings. Before and during recordings, all surface areas—including manipulators, microscope knobs, computer keyboard, etc.—that the experimenter needed to contact during experiments were cleaned with RNase Away solution (Molecular BioProducts). After recordings, the cell’s cytosol was aspirated via the glass pipette used for recording. Although the aspirated cytosol may have contained genomic DNA, our choice of cDNA preparation, which involved poly-A based mRNA selection, eliminates the possibility of genomic contamination in the RNAseq data. For sample collection, we quickly removed the pipette holder from amplifier head stage and used positive pressure to expel samples into microtubes containing cell collection buffer while gently breaking the glass pipette tip. Cell collection microtubes were stored on ice until they were used.

#### * cDNA Library Preparation

Same procedures were followed as described in Földy et al. (2016) and Winterer et al., (2019). Single-cell mRNA was processed using Clontech’s SMARTer Ultra Low RNA Input v4 or SMART-Seq HT kit. As first step, cells were collected via pipette aspiration into 1.1 µL of 10x collection buffer and spun briefly before they were snap frozen on dry ice. Samples are stored at −80 °C until further processing, which was performed according to manufacturer’s protocol. Resulting cDNA was analyzed on the Fragment Analyzer (Advanced Analytical). Library preparation was performed using the Nextera XT DNA Sample Preparation Kit (Illumina) according to manufacturer’s protocol. Following library preparation, cells were pooled and sequenced using NextSeq 300 high-output kit in an Illumina NextSeq 500 System with 2×75 paired-end reads.

### Bioinformatics

#### * Online Available RNA Sequencing Data

Original fastq files containing raw reads for GSE99888 (Harris et al., 2018) were downloaded from NCBI GEO (www.ncbi.nlm.nih.gov/geo/GSE99888).

#### * Processing of RNA Sequencing Data

After sequencing, raw sequencing reads were aligned to the Ensembl GRCm38 reference transcriptome (Version 95), using Kallisto’s *quant* command (Bray et al., 2016) with 100 bootstraps. For convenience, Ensembl gene IDs were converted to gene symbols using a reference file generated by biomaRt (Durinck et al., 2009). In the few cases where different Ensembl gene IDs identified the same gene symbol, transcript per million (TPM) levels were summed.

#### * Quality control

All data analysis was performed using R and Python codes. First, in each cell, we calculated the number of unique genes and the number of aligned reads. Second, we calculated the median and median absolute deviation of these two values across all cells. Cells that had either value more than 3 median absolute deviations below the median were removed as failing quality control.

#### * Normalization of Gene Expression

Transcript per Million (TPM) normalization of transcripts was calculated by a built-in Kallisto function.

#### * Differential Gene Expression

For calculating differentially expressed (DE) genes, we first read in Kallisto’s output using Tximport (Soneson et al., 2015), to account for uncertainty in alignment. We then imported the results to edgeR and used a quasi-likelihood test on all genes that were expressed in at least 5 cells in the two groups being compared. Genes were labeled as DE if there was a fold difference of at least 2 (absolute value of logFC>1) in average expression, at a significance of *p*-adjusted<0.05.

#### * Dimension Reduction Methods and Analysis

To plot high dimensional data, we used five dimension reduction algorithms: Principle Component Analysis (PCA), t-distributed stochastic neighbor embedding (t-SNE), Fast Fourier Transform-accelerated Interpolation-based t-SNE (FIt-SNE), negative binomial t-SNE (nbt-SNE), and Uniform Manifold Approximation and Projection (UMAP). All methods transform high dimensional data to a lower dimension while preserving key information. PCA is a linear transformation which attempts to preserve the variance in the positions of cells. t-SNE is a non-linear transformation that attempts to preserve the distances of cells only to their nearest neighbors, losing macro-scale information in the process. Both FIt-SNE and nbt-SNE are modifications to t-SNE. FIt-SNE is designed to make t-SNE run faster on large scale data and attempts to preserve macro-scale information, while nbt-SNE modifies the distance function used from a Gaussian to a Negative Binomial model that is believed to be more accurate for RNA-sequencing data. UMAP works similarly to t-SNE, but uses a modified distance function and attempts to preserve macro-scale information by putting more weight on distances between farther away points. While PCA is unable to capture more complex, non-linear information, it has the advantage of interpretability; only on a PCA plot do the distances along each axis have any biological meaning.

#### * Classification Accuracy

Related to Figs. 1 and 6. To determine how accurately cell types could be classified, we trained a Random Forest Classifier algorithm. Briefly, a Random Forest creates multiple decision trees to classify cells, using only different subset of the genes each time. Each of the individual trees ‘votes’ on a classification, and the most popular classification is used. We used an ensemble of 100 decision trees for our algorithm. To not bias the results of the Random Forest, and to simplify interpretation, we randomly removed cells from the larger class so that the two categories had the same number of cells. This allowed us to label 50% accuracy as the base line of what we would get if there were absolutely no differences between the two categories. We then used 80% of the cells as a training set and evaluated the result on the remaining 20%. For each classification we repeated this method 100 times, each time randomly selecting which cells were removed, and which were used in the training and test set. This allowed us to get an average classification accuracy for the categories as a whole, as well as for each cell.

#### * Gene Selection

Related to Fig 2. In a bioinformatics data set, when exploring the difference between multiple cell types, or trying to identify cell types via clustering, most genes do not contribute any information to the separation. Continued consideration of these genes can decrease the signal to noise ratio to the point where existing distinctions cannot be resolved. As such, it is important to first trim the list of genes to only keep significant genes. However, there is no singular ‘best’ approach to select these genes for specific types of problems, let alone all bioinformatics analysis in general. As such, we tried a number of different gene selection methods to confirm if any of them would give a clear separation. Three of them (chi-squared, mutual information and ANOVA F-value) were already implemented in *sklearn*. For chi-squared we used log2 of gene TPM expression levels, while for mutual information and ANOVA F-value we used a boolean value of whether or not a gene was expressed. In all three cases, we took the best separating 150 genes. Next, we tried the 150 genes that were used for classifying the CA1-IN data. We also tried the top 150 genes that correlated with these separators but were not them. Lastly, we ran a method described by Kobak et al. (2019) that finds the most relevant variable genes accounting for expression rate, and using a cut-off of TPM>32, and once again took the top 150 genes.

#### * Transcriptomic Mapping

Related to Figs 2 and 4. To map our data onto the CA1-IN data set, we used the method described in Kobak et al. (2019). Briefly, after key genes were selected, we used the correlation coefficient to find the k-nearest neighbours for each cell. We took the median position of the embeddings of these k neighbors, as the mapping position of our cells.

#### * Sliding Window

Related to Fig. 7. To determine whether the expression of gene was ‘upregulated’ or ‘downregulated’ at a given age, we only considered gene expression data in a binary format, i.e. expressed or not in a single cell. Since Kallisto’s bootleg approach has shown to occasionally assign very low expression levels to transcripts that are not expressed, we used a low cut-off (0.6 TPM) for determining if a gene was expressed or not. To increase the statistical power, we ignored genes that were expressed in less than 6 cells, or not expressed in less than 6 cells. After that, for each gene we used a sliding window to calculate the transition point with the highest loss of Gini impurity. To calculate the p-values, we used a Monte Carlo simulation. For each potential number of cells that a gene might be expressed in, we ran 100,000 simulations by randomizing the expressions and calculated the Gini Impurity loss for each. We then used these distributions to calculate the p-value for each gene.

#### * Gini Impurity

Related to Fig. 7. Gini impurity is a measure of the degree of heterogeneity of a group. If elements in a set were randomly labeled based on the distribution of categories in a set, the fraction that would be incorrectly labeled is the Gini impurity. It can be calculated as 1 - Sum(p_i_), where p_i_ are the fractions of a set that belong to each group. The number varies from 0 (complete homogeneity) to almost 1 (every element is in a different group), and from 0 to 0.5 when there are only two groups. If a set is divided into two smaller parts, the average Gini impurity - normalized by the sizes of the two sub-sets - of the two sub-sets will be smaller than that of the entire set. This difference is a measure of the information gain from the separation.

#### * Linear Support Vector Machine with Recursive Feature Elimination

Related to Figure S4. Support vector machines are a classification algorithm. They find the best line (2 features), plane (3 features), or hyperplane (4 features and above) along which to separate the data for classification. Linear support vector machines use features as is, rather than generating new features, making them simple to interpret. Recursive feature elimination is a way to reduce the number of features used in a classification algorithm. Briefly, after a classification algorithm is trained, the features are weighted and the least important features are dropped. These two steps are repeated until the dataset is reduced to a previously selected number of features. For our data, genes were the features, and cell types were the classes that we tried to separate. Using the two algorithms together, we found the top 50 genes that best classified cell types using a linear support vector machine.

## Acknowledgments

We thank Drs. János Szabadics and Jochen Winterer for discussions. This work was supported by funding from the Swiss National Science Foundation (Switzerland, CRETP3_166815 and 31003A_170085) and the Dr. Eric Slack-Gyr-Stiftung (Switzerland) to C.F.

## Author contributions

L.Q. performed electrophysiological, morphological and single-cell RNA-seq experiments. D.L. performed bioinformatic analyses. C.F. wrote the manuscript. C.F., D.L, L.Q. designed the study, analyzed data, prepared figures, and edited the manuscript.

## Data sharing

GSE142546

## Supplementary Information

**Fig. S1.**
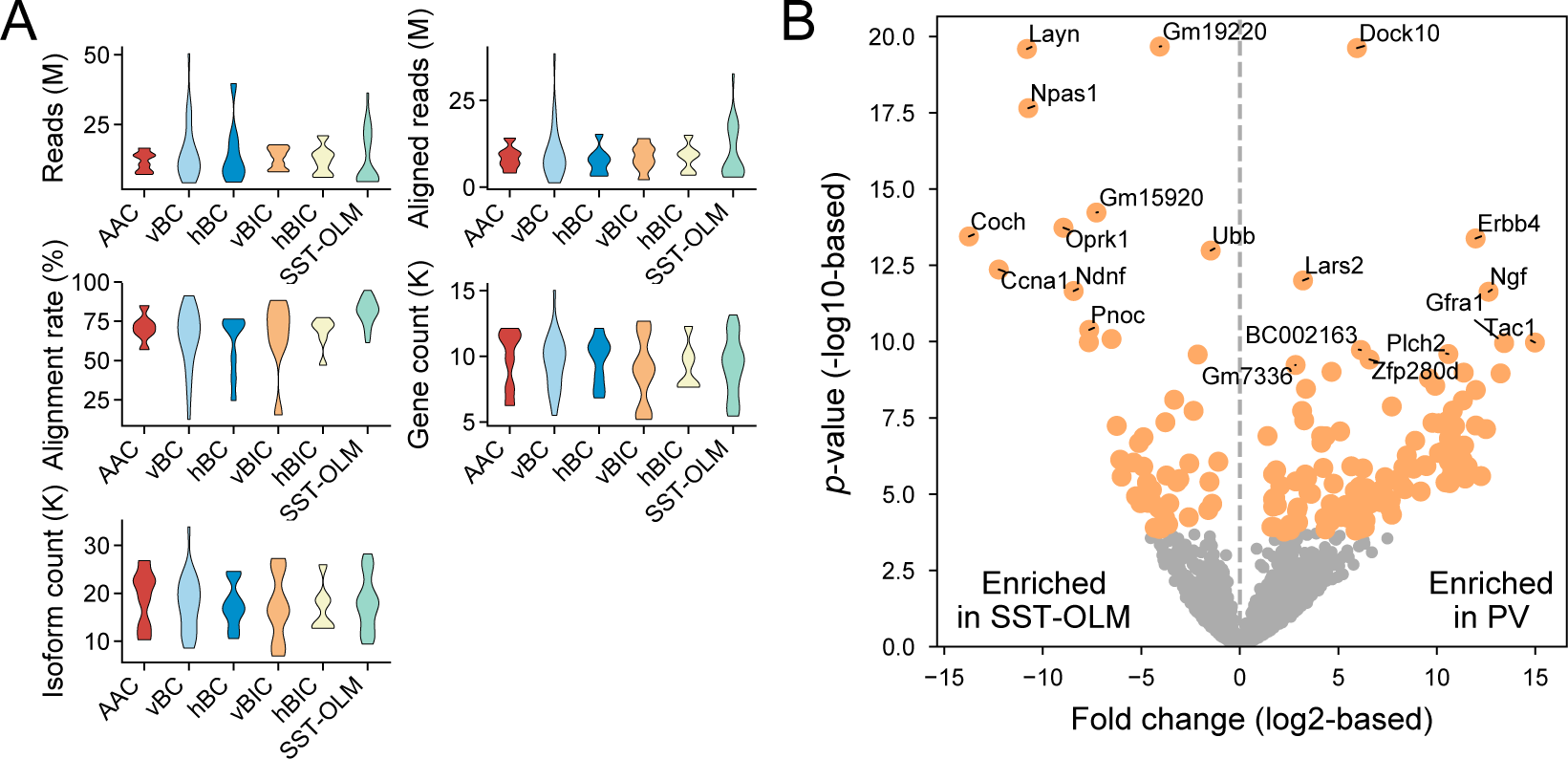
Single-cell RNAseq profiling of PV-INs. We first made comparisons between this current PV set to a previously generated SST interneuron data set. We hypothesized that transcriptomic differences between PV
and SST cells would define an ‘upper limit’ of molecular distinctions that may be present between the different morphological PV types. **(A)** Violin plots show quality control parameters (number of reads, aligned reads, alignment rate, gene count, and isoform count) after single-cell sequencing for 5 morphological PV and for the SST-OLM type, which data set was adopted from Winterer et al. (2019) and generated with identical methods
as the PV set of this study. **(B)** Before examining differential gene expression between morphological PV types in detail, we analyzed the major differences between PV and SST cells. Volcano plot shows differential gene
expression between PV, where all morphological types were pooled together, and SST-OLM cells. Each circle
represents a single gene. Orange color denotes genes that are differentially expressed between the two cell types
with a 2-fold difference (|log2|>1) and an adjusted p<0.01. Statistics were calculated using *edgeR*. The number of differentially enriched genes are n=47 and 124 in SST-OLM and PV-INs, respectively.

**Fig. S2.**
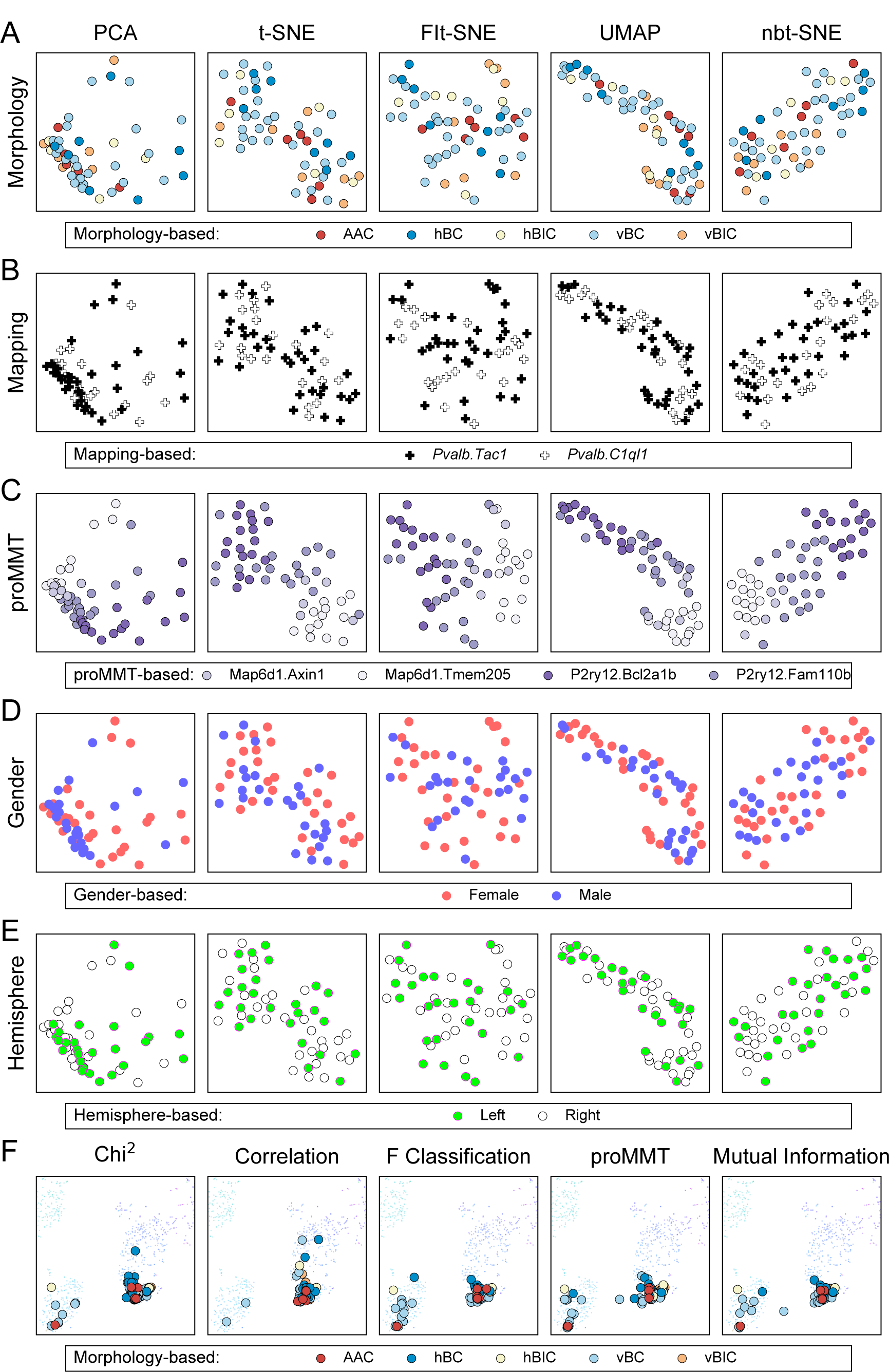
Transcriptomic characterization of PV-INs. **(A-E)** Our transcriptomic characterization of PV-INs using proMMT revealed four clusters, but visualization using nbt-SNE could not clearly separate these clusters in the two-dimensional space (Fig. 2A). Here, using multiple different visualization approaches, we show invariant similarly and lack of separation in transcriptome-based clustering of PV-INs. The five visualization approaches that we used are PCA, t-SNE, Flt-SNE, UMAP, and nbt-SNE, and are labeled on the top of each column. Note that since all plots in a column use the same visualization technique, the location of symbols (representing single cells) are the same. However, in each row, we colored the symbols differently, according to different classification scenarios that we considered in addition to their proMMT-derived cluster. Cells are labeled according to the 5 morphological PV types. None of the five visualization techniques revealed additional separation in the two-dimensional space than nbt-SNE (panel A). Cells are labeled according to whether they mapped onto *Pvalb.Tac1* or *Pvalb.C1ql1* (panel B; see Fig. 2B and C). Cells are labeled according to their proMMT clusters (panel C; the rightmost nbt-SNE plot is shown in Fig. 2A). Cells are labeled according to the gender of the animal from which they were collected from (panel D). Cells are labeled according to the side of the hemisphere they were collected from (left or right hemisphere). To conclude, these visualizations show a lack of transcriptomic separation based on morphological, gender, or hemispherical properties (panel E). **(F)** Plots show mapping of our PV-INs onto the PV-INs of the CA1-IN data set, using 5 different methods of feature selection; chi-squared, F Classification and Mutual Information were used on the PV-INs from the CA1-INs data set to find the best markers for *Pvalb.Tac1* and *Pvalb.C1ql1* clusters in that data set. proMMT represents clustering based on the 75 genes selected by the algorithm, and Correlation is based on the top 150 genes that best correlated with the 150 gene used in the CA1-IN study, but not including those (see also Fig. 2C).

**Fig. S3.**
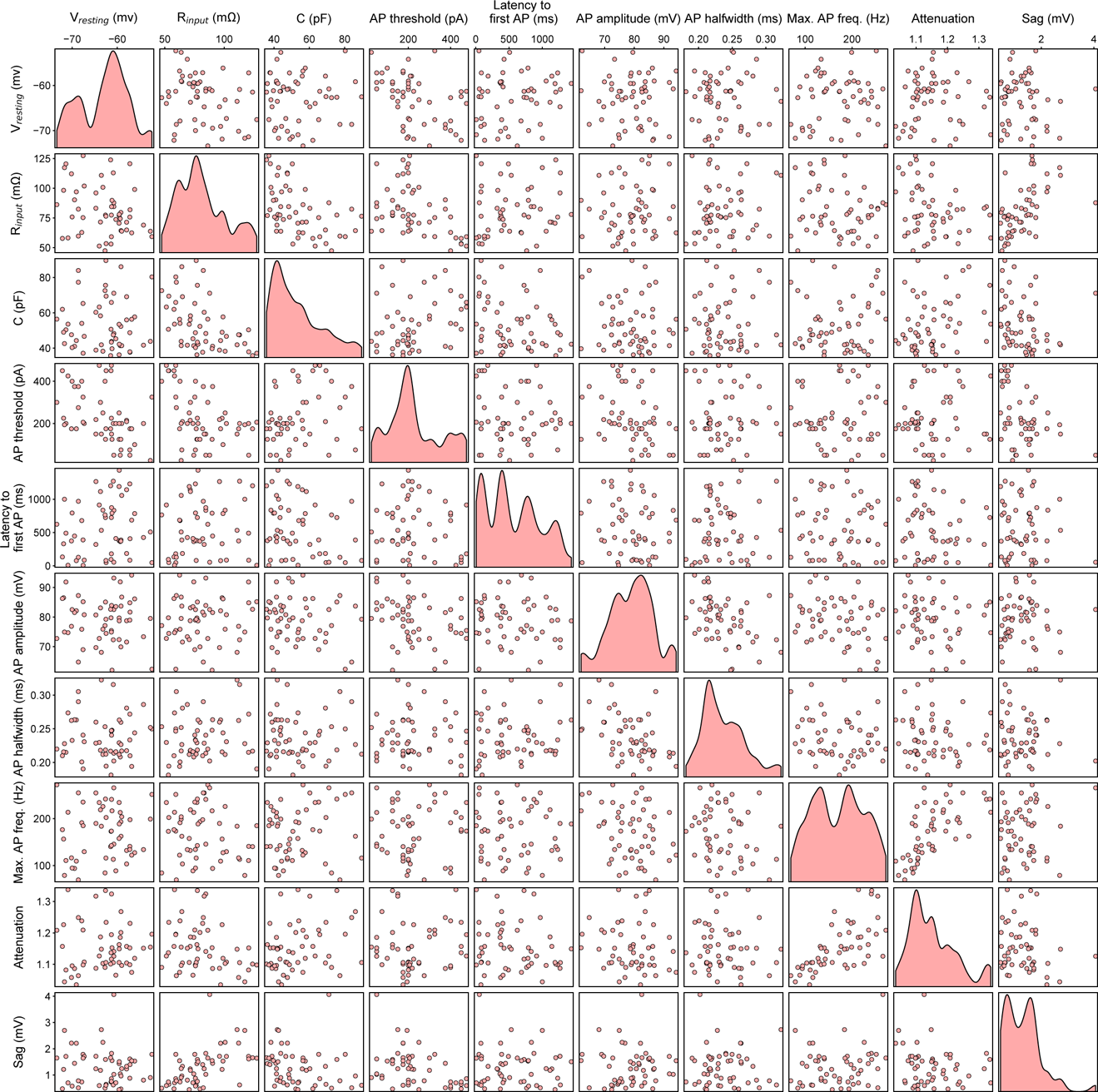
Cross-comparison of electrophysiological parameters measured in PV-INs. In Fig. 3, we show that unbiased clustering of electrophysiological parameters measured in PV-INs did not reveal biophysical PV types. To support this conclusion, here we considered the possibility that hidden pair-wise correlations between different electrophysiological parameters may define distinct biophysical clusters, not detectable by unbiased clustering. To detect these, we made scatter plots displaying data points of each pair of ten electrophysiological parameters we analyzed. However, none of these plots revealed distinctly separating clusters, indicative of different biophysical types among hippocampal PV-INs. Plots along the diameter display histograms showing the distribution of the corresponding parameter measured in all >P21 PV-INs (n=53) for which electrophysiological measurements were available, independent of cells’ morphological and transcriptomic type.

**Fig. S4.**
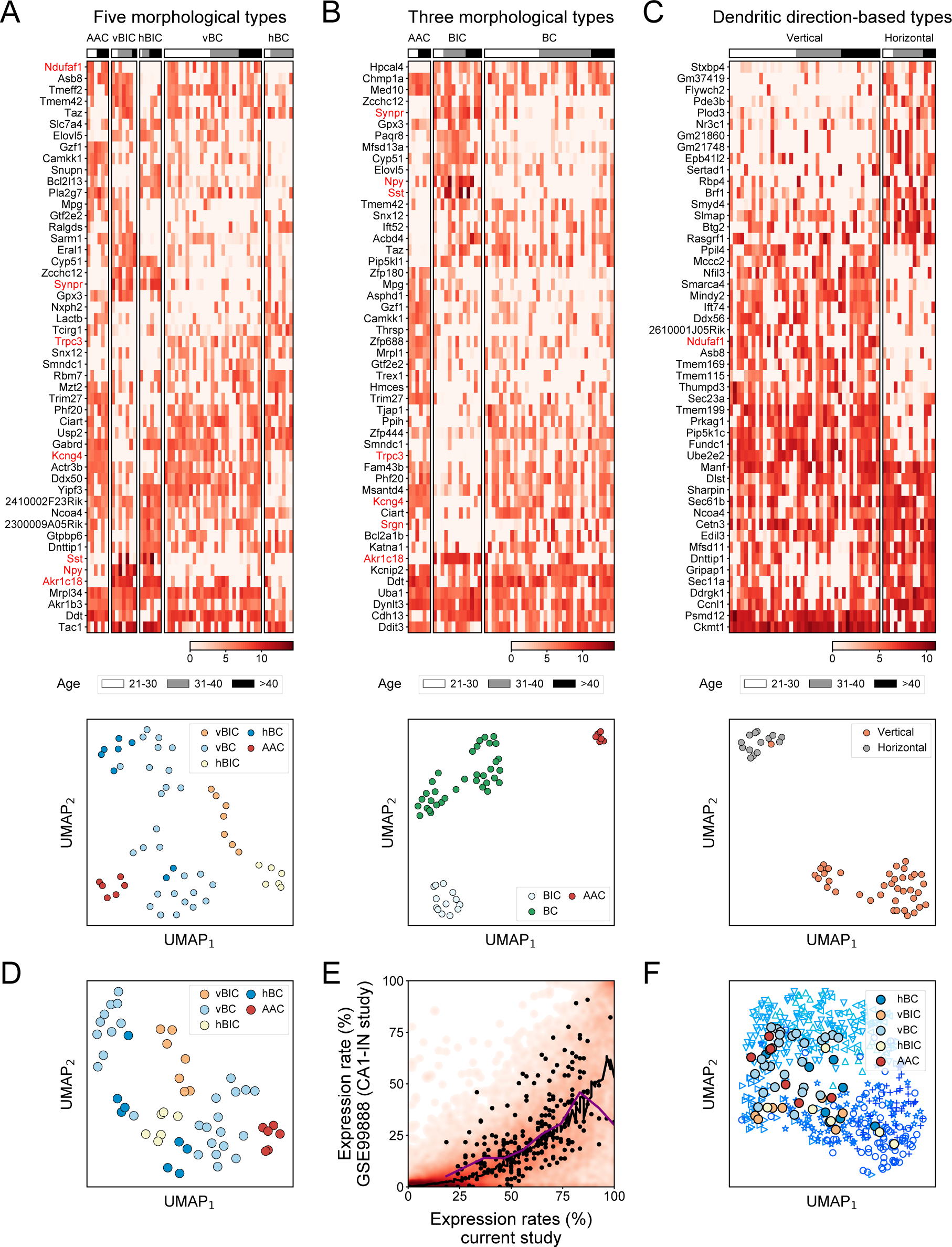
Support vector machine classification and gene selection in morphological PV types. In Fig. 4, we used statistical comparisons to identify enriched genes among different morphological PV types. Here, we implemented random forest classification to this problem. (A-C) Heat maps (upper plots) of gene sets identified by linear support vector machine (svm) classification between the 5 morphological PV types (recursive feature elimination was used to select 50 genes; panel A), axo-morphological types (panel B), and dendro-morphological types (panel C). Cells are ordered according to age. Note that individual genes are not necessarily significantly different between any two morphological type. Genes names colored in red are statistically significant (using *edgeR*, fold difference>2, p<0.05) and are shown in Fig. 4. UMAP plots (lower plots) were made using all genes in their respective heat map. These highlight clear separation between each of the morphological types. **(D)** UMAP plot of cells, colored by the 5 morphological PV types, using an extended set of n=124 genes from A-C. **(E)** Comparison of expression rate of genes in PV-INs from this current versus the CA1-IN study, displayed as a heat map. Black line marks loess regression fit. Black points label the 124 svm genes from A-C. Black line marks loess regression fit of all genes, and purple line marks loess regression fit of svm genes. Contrary to the differentially expressed genes in Fig. 4, selected genes followed the expression pattern of all genes (black line), indicating that they did not experience unexpected dropouts in the CA1-IN data set. **(F)** UMAP based embedding of PV-INs from the CA1-IN study (Harris et al., 2018), and mapping the PVINs of this current study onto the UMAP embedding using the 124 svm genes for A-C. Symbols refer to the following transcriptomic subtypes, as described in the original study: Δ *Pvalb.Tac1.Akr1c18*, Δ *Pvalb.Clql1.Cpne5*, Δ *Pvalb.C1ql1.Npy*,Δ*Pvalb.Clql1.Pvalb*, + *Pvalb.Tac1.Nr4a2*, ○*Pvalb.Tac1.Sst*, and ★ *Pvalb.Tac1.Syt2*. In conclusion, while for technical reasons the random forest approach revealed a largely different set of genes than pair-wise statistical comparisons between the morphological types, these outcomes corroborate our finding that specific gene sets identify morphological PV types.

**Fig. S5.**
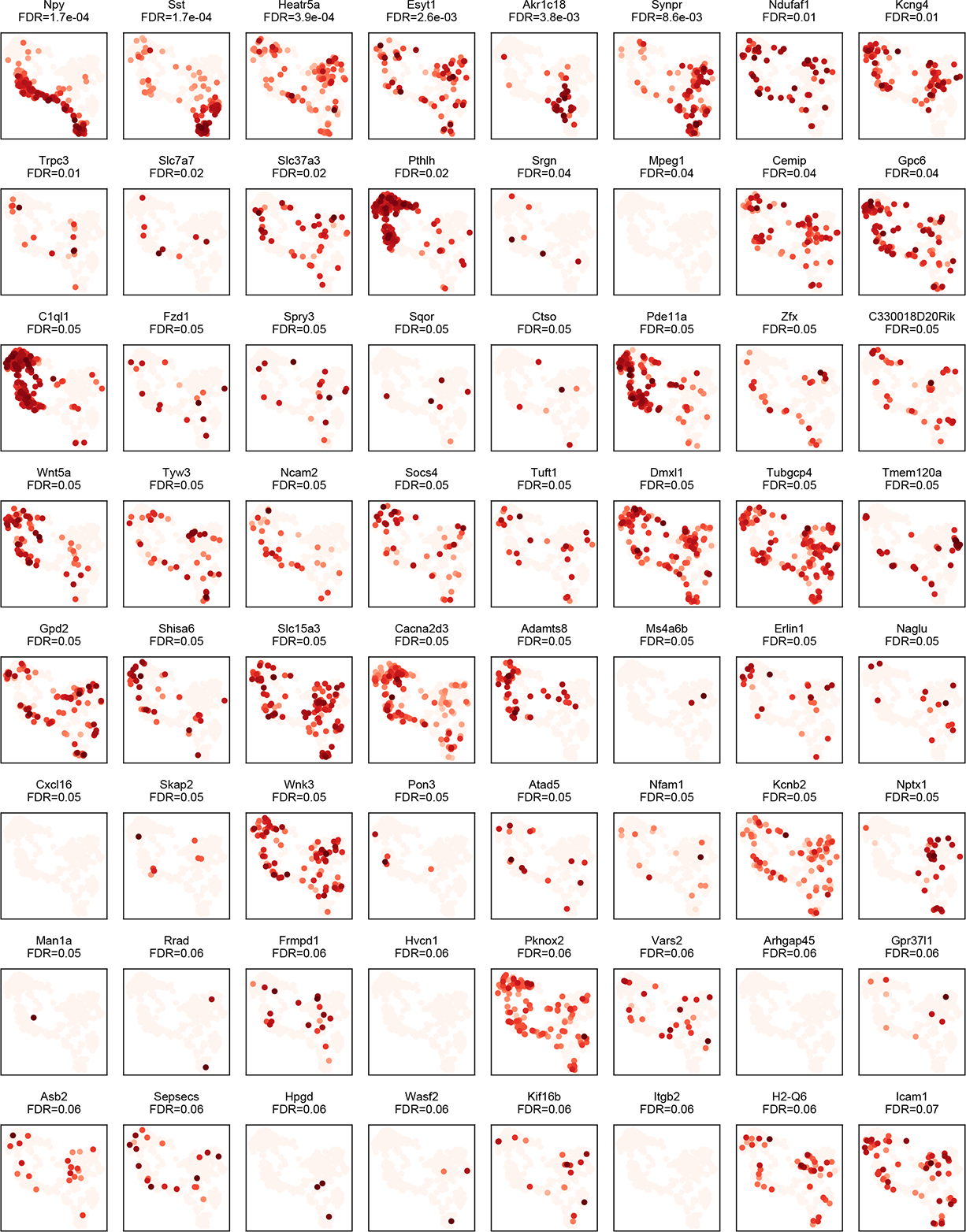
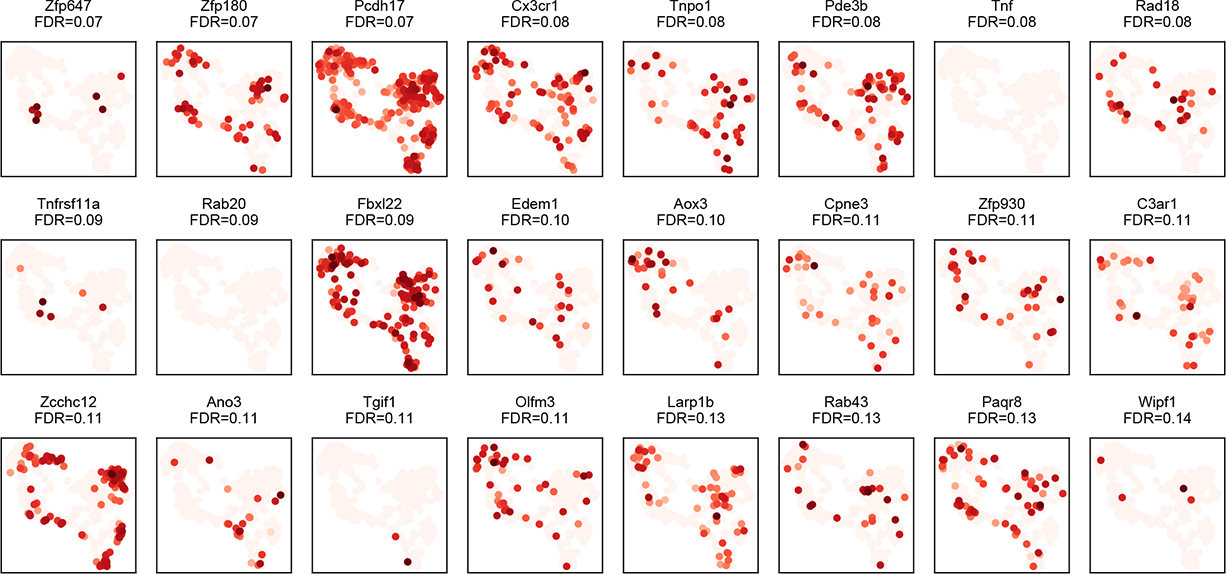
Expression of morphology-associated genes in the CA1-IN data set (Harris et al., 2018). Using a morphology-based supervised approach, in Fig. 4, we identified a set of n=88 genes, which separately clustered BIC and non-BIC cells, and outlined further distinctions among the 5 morphological PV types. (A) Using the same UMAP representation as shown in Fig. 4F, we show the expression of each of the morphology-associated 88 genes in the CA1-IN data (each plot is labeled on top with the corresponding gene name). Together, these plots highlight that 7/88 genes were not at all, and further 25/88 genes were detected only in ≮20 of the n=479 PV-INs. As one example, *Trpc3*, which we found to be differentially expressed between BIC and non-BIC cells (see also Lee et al., 2011), was expressed only in 18/479 cells (3.7%). By contrast, we detected *Trpc3* in 24/54 cells (44.4%; for completeness, this count also includes BIC-s, which do not express this gene) in our data set. While the two data set were generated with different approaches (e.g. patch-seq versus FACS-based cell collection), currently there is no cause known for this discrepancy. However, our gene-by-gene analysis highlights a limitation to which the morphology-associated genes identified in this current study could infer the morphological identity of PV-INs in the previous CA1-IN data set.

**Fig. S6.**
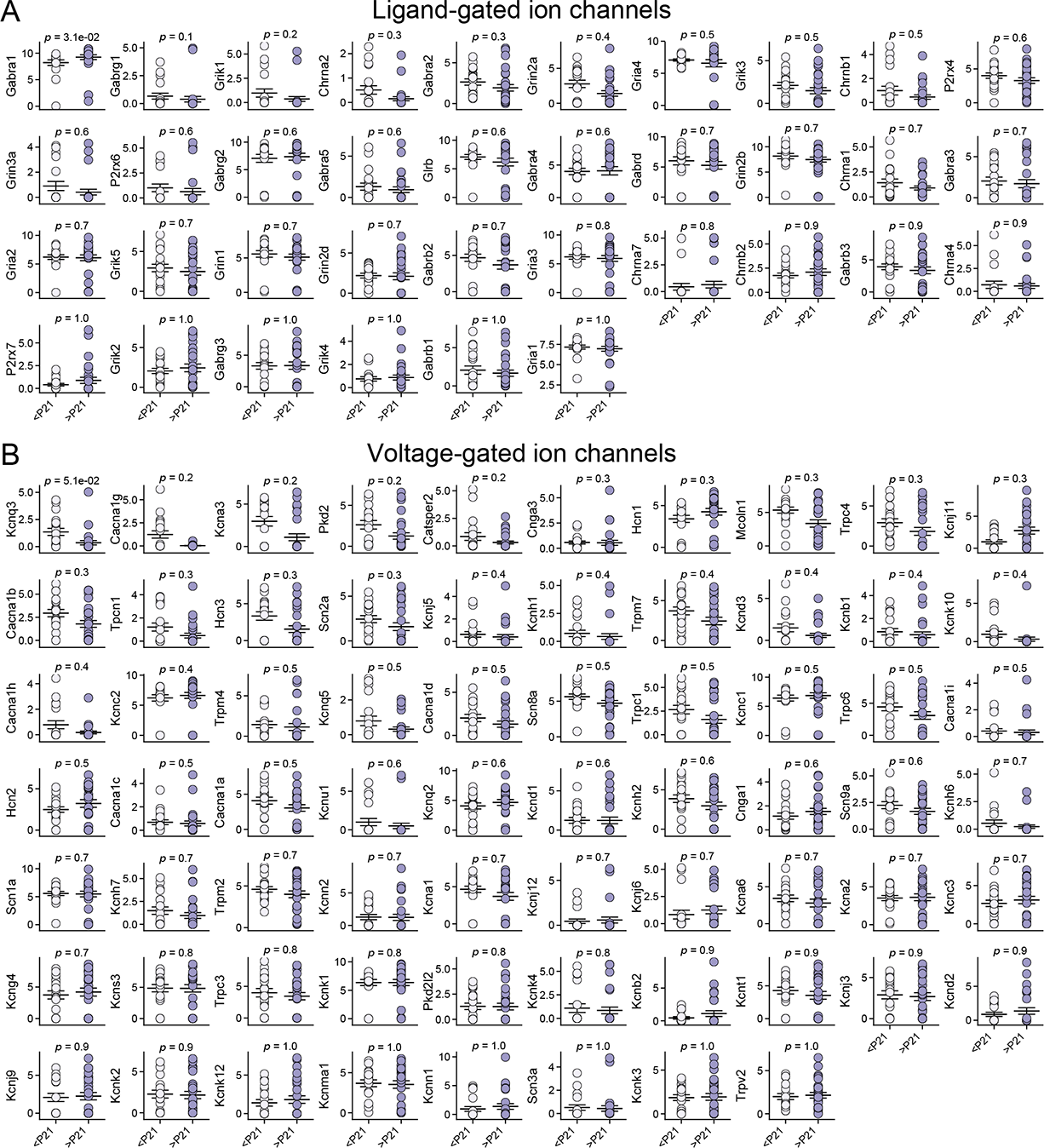
Morphological and electrophysiological analysis of vBC type PV-INs during circuit maturation. In Fig. 6, we identified physiological features in BC-s, which correlate with age. Here, we show further evidence supporting these findings. (A) Plots show pair-wise comparison of ligand-gated ion channel expression in <P21 versus >P21 vBC type cells. Each plot is labeled with the name of the corresponding gene. Ordinate values represent normalized log2(TPM) gene expression level. Abscissa values are labeled only in the bottom, but applicable to all plots. The order of the plots was determined by the *p* value of statistical significance (two-sided Mann-Whitney test). **(B)** Plots show pair-wise comparison of voltage-gated ion channel expression in <P21 versus >P21 vBC type cells.

**Fig. S7.**
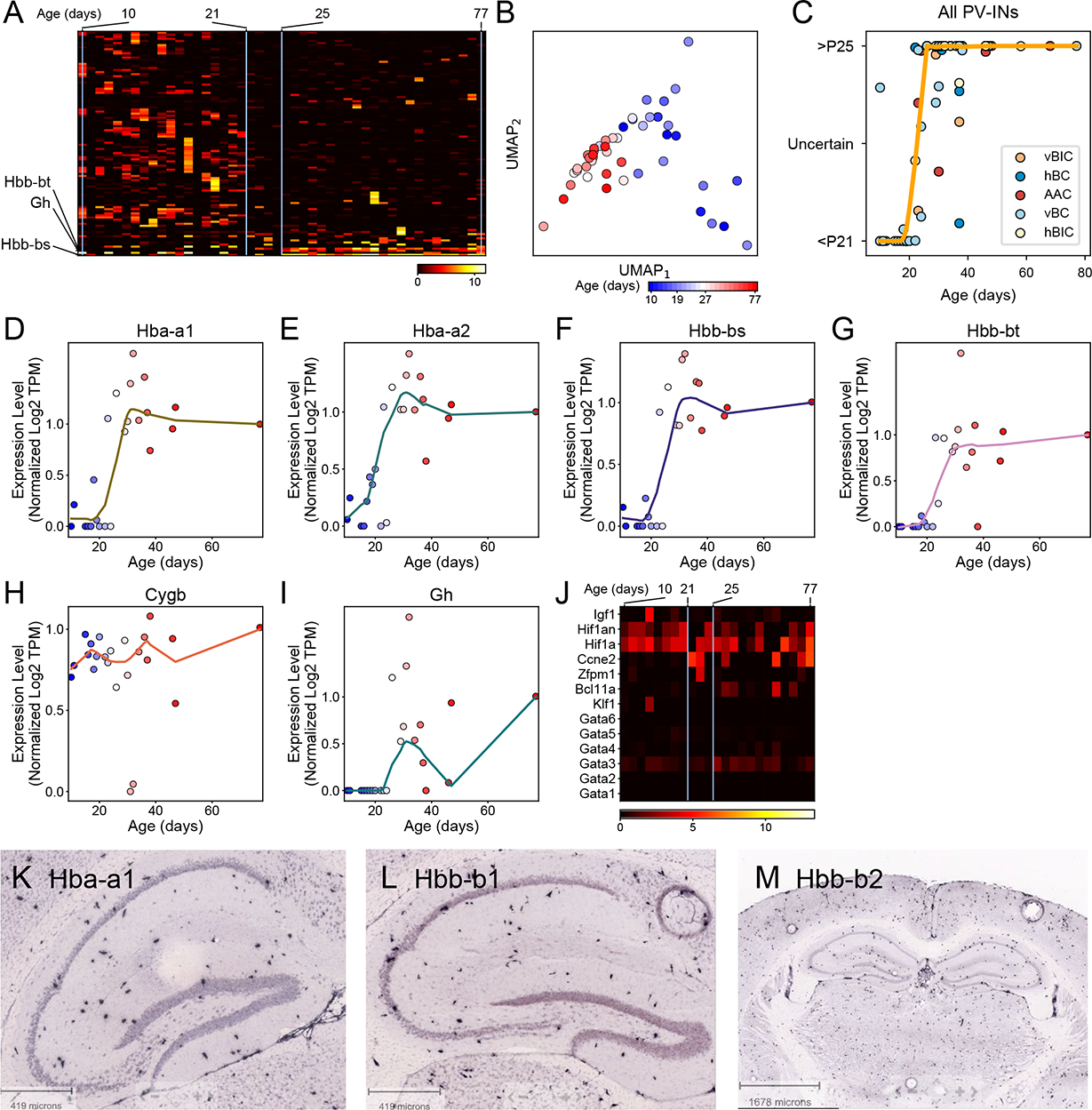
Hemoglobin-related gene expression changes in PV-INs. In Fig. 7., we identified rapid and transcriptomic changes and an age-dependent onset of hemoglobin expression in PV-INs. Here, we provide more detailed evidences supporting this finding. **(A)** Heat plot shows the expression level of genes identified by Monocle in vBC type cells. GO analysis revealed the presence of multiple gene families, none of which were statistically significant (not shown). Cells (columns) are ordered according to age. **(B)** Using the genes identified in panel A, UMAP shows age-dependent separation of vBC type cells. **(C)** Plot of consistency with which a PVIN cell’s age can be predicted using Random Forest Classifier based on genes from Fig. 7 panel A, fitted with a loess curve. **(D-I)** Normalized loess fits of growth hormone (Gh) and different globin gene expression levels versus cell age in vBC type cells. **(J)** Heatmap shows the expression of Hb gene regulating factors in vBC type cells, which were sorted by age (left to right). Cells (columns) are ordered according to age. **(K-M)** Photographs show hippocampal ISH data adopted from Allen Mouse Brain Atlas of Hb subunits Hba-a1 (panel K), Hbb-b1 (L), Hbb-b2 (M), respectively. Sparsity of expression and localization of cell bodies suggest that Hb expression is feasibly restricted to PV-INs. All samples are from P56 male mice (C57BL/6J).

